# Lung-targeted cytokine-coding RNA-lipoplexes induce T and NK cell-mediated anti-tumor immune response

**DOI:** 10.64898/2026.05.06.723126

**Authors:** Ayline Kübler, Sophie-Christin Linkenbach, Fulvia Vascotto, Elif Diken, Özlem Akilli, Eliana Stanganello, Aniello Federico, Sina Fellermeier-Kopf, Alexander Muik, Friederike Gieseke, Martin Suchan, Ferdia Bates, Kaushik Thanki, Hossam Hefesha, Isaac Hernan Esparza Borquez, Matthias M. Gaida, Jutta Petschenka, Kerstin C. Walzer, Jürgen Brück, Matthias Miederer, Sebastian Kreiter, Mustafa Diken, Ugur Sahin

**Author notes:** Corresponding authors. (U.S.), (A.K.), mailing address: TRON gGmbH, Freiligrathstr. 12, 55131 Mainz, Germany. Boehringer Ingelheim International GmbH, Binger Straße 173, 55216 Ingelheim am Rhein, Germany (Jutta Petschenka). Department of Translational Imaging in Oncology, National Center for Tumor Diseases (NCT/UCC) Dresden: Faculty of Medicine and University Hospital Carl Gustav Carus, University of Technology Dresden (TUD), Dresden, Germany, German Cancer Research Center (DKFZ) Heidelberg, Germany, and Helmholtz-Zentrum Dresden-Rossendorf (HZDR), Dresden, Germany.

## Abstract

Lung is a major site of metastases for many primary cancers associated with poor outcomes. A central challenge in cancer immunotherapy is overcoming tumor immune evasion, which limits effective antitumor responses. Here, we investigated whether combinatorial mRNA-encoded cytokine therapy can overcome tumor immune evasion by coordinately engaging innate and adaptive immunity, using murine models of pulmonary metastases. We employed intravenously administered cationic nucleoside-modified mRNA-lipoplexes (RNA-LPX) for targeted delivery of mRNA-encoded cytokines to the lung. The cytokine mix containing interferon-α, half-life extended interleukin (IL)-7, and a half-life extended IL-2 variant with reduced CD25-binding modulated the tumor immune microenvironment resulting in a potent and broad anti-tumor response and prolonged survival with good tolerability at the conditions tested. Using cell depletion experiments, we demonstrated that both T and natural killer (NK) cells are crucial mediators of the observed anti-tumor efficacy of the cytokine RNA mix, which induced activation and effector function of NK and T cells, coupled with reduced regulatory T cells (T_reg_) numbers and T_reg_ activation in the lung. Importantly, antitumor efficacy was maintained in models of impaired antigen presentation, including loss of an immunodominant tumor antigen and MHC class I deficiency, where NK cells served as the primary effectors. The cytokine RNA mix induced immune cell activation in the primary human lung tumor culture, suggesting potential for translational application. Collectively, these findings demonstrate that combinatorial cytokine therapy can drive both antigen-dependent and antigen-independent tumor control for the treatment of lung metastases.

## MAIN

Lung is one of the most frequent sites of metastases for several cancer types, such as lung cancer, colorectal cancer or, melanoma [1]. The presence of distant metastases has a profound negative impact on prognosis: e.g. in colorectal cancer the occurrence of lung or liver metastases is associated with a drop in the 5-year survival from 91% to below 10% [2], increasing the need for new innovative treatment strategies. Cytokines have recently gained attention as potential anti-cancer immunotherapy agents. IL-2 is crucial for T cell homeostasis and expansion and has potent ability to stimulate cytotoxic T cells and NK cells [3]. IL-7 mediates the homeostasis of both memory and naïve T cell populations and is primarily important for survival of memory CD8^+^ T cells [4]. The anti-tumor activity of interferons has been recognized for at least half a century. Interferon-α (IFN-α) can mediate direct anti-tumor effects as well as dendritic cell (DC), NK and innate lymphoid cell (ILC) activation, increased antigen presentation, induction of secondary mediators and inhibition of regulatory T cells (T_reg_) and myeloid-derived suppressor cells (MDSC)[5]. The US FDA approved recombinant IL-2 and IFN-α as immunotherapy for several malignant diseases [6] which marked a milestone in cancer immunotherapy as clinical proof of boosting the endogenous anti-tumor immune response. However, low response rate, short serum half-life of IL-2 and high toxicity associated with high-dose IL-2 and IFN-α administration limited their clinical use [6] and led to the development of next-generation IL-2 variants and strategies to increase half-life, as for example albumin fusion [7].

In this study, we utilized cationic RNA lipoplex (RNA-LPX) particles that specifically target the lung [8–10] to investigate the anti-tumor activity of RNA-encoded engineered cytokines in a mouse model of lung metastases. We used a mix of an albuminated IL-2 variant with reduced CD25 binding (Alb-IL-2mut), an albumin fusion of IL-7 (Alb-IL-7) and IFN-α to modulate the tumor immune microenvironment (TIME) in experimental models of lung metastases. We show that this approach activates both the innate and adaptive immune response, resulting in anti-tumor activity even in immune escape models (gp70 or β2 microglobulin knockout), extended survival of mice, and protection against tumor re-challenge. Our results demonstrate feasibility of targeted delivery of RNA-encoded cytokines and cytokine variants to the lung and provide mechanistic insights into immune-mediated suppression of lung metastases.

## RESULTS

### Delivery of RNA-encoded cytokines to the lung

We previously reported that RNA-LPX containing equal parts of DOTMA and cholesterol at a cationic lipid to RNA ratio of 4:1 specifically target the lung [8,9]. To improve and prolong protein translation and to reduce the mRNA stimulatory capacity via innate receptors, we incorporated N1-methylpseudouridine triphosphate (m1ΨTP) instead of uridine 5′-triphosphate into the RNA [11]. Upon intravenous (i.v.) administration of m1Ψ-modified luciferase (*luc*) RNA-containing LPX to non-tumor bearing mice, we confirmed luciferase expression in the lungs as shown by bioluminescence imaging (BLI; Fig. 1A-C). Histological stainings revealed dominant targeting of CD31^+^ lung endothelial cells (Supplementary Fig. S1A, B). RNAscope and histological analysis of CT26-tumor-bearing lungs showed accumulation of RNA around blood vessels 1 hour after injection (Fig. 1D), followed by protein expression allover the lung as well as in tumor nodules 6 and 24 hours after injection (Fig. 1D, Supplementary Fig. S1C, D).

**Figure 1:**
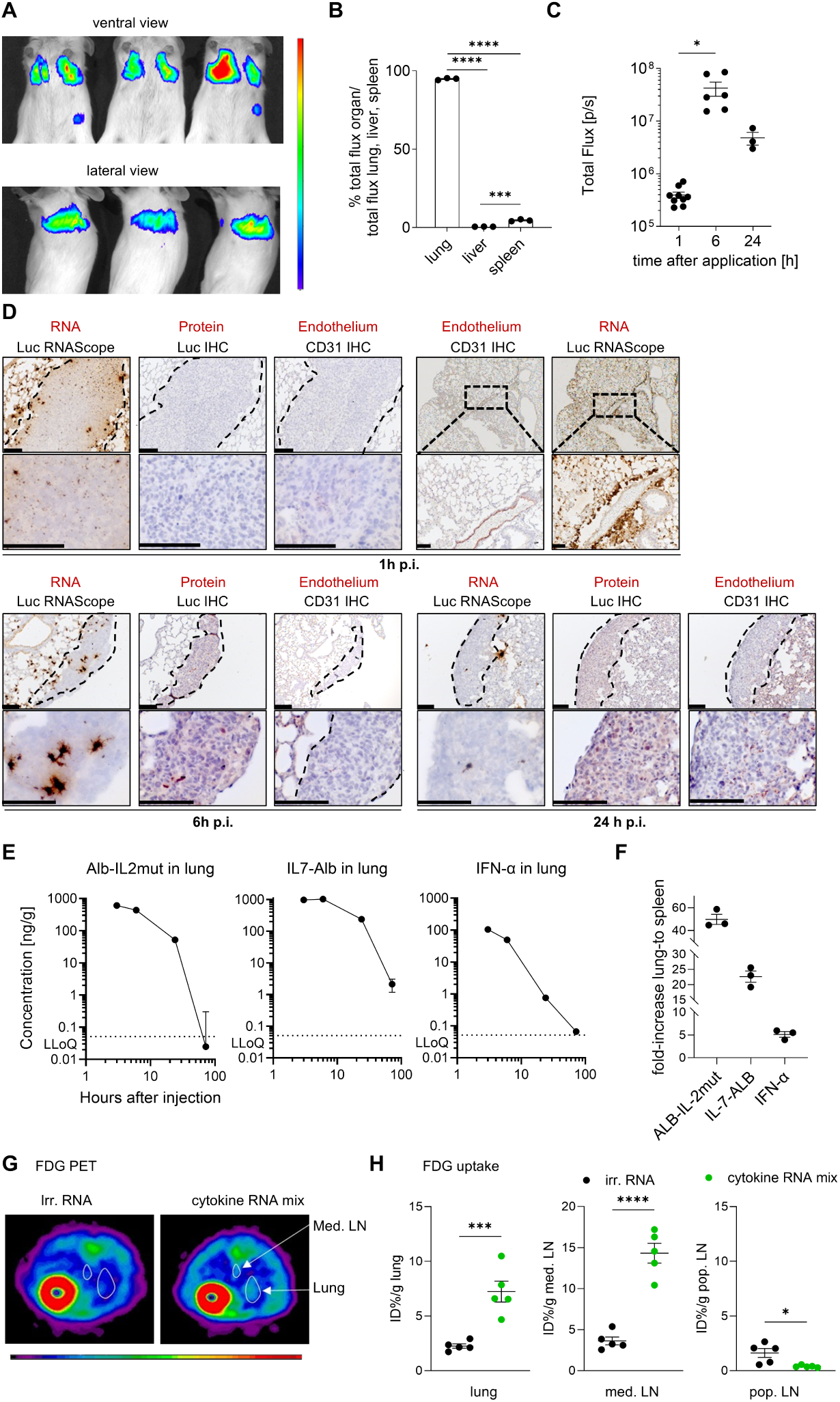
Intravenous administration of RNA-LPX delivers RNA-encoded cytokines to the lung. (**A-D**) Luc–encoding RNA formulated as RNA-LPX was administered i.v. to non-tumor bearing (A, B; n=3) or CT26 metastases-bearing (C, D) BALB/c mice (n=3-9). (**A**) BLI of total body, (**B**) normalized organ signal from explanted organs. Data were analyzed by ordinary one-way ANOVA and Tukey’s test for multiple comparisons. (**C**) Kinetics of BLI of the lung signal in metastases-bearing mice, significance was determined by mixed-effects analysis with Geisser-Greenhouse correction and Tukey’s test for multiple comparisons. (**D**) *Luc* RNA, luc protein and CD31 (PECAM-1) were detected via RNAscope and immunohistochemistry on consecutive sections 1, 6 or 24 hours post injection, scale bar: 100 µm, tumor tissue is indicated with dashed line. (**E**) Cytokine quantification in lung after the indicated time points. (**F**) Cytokine fold increase in lung normalized to spleen at 6 hours after injection. (**G**) FDG positron emission tomography (PET) imaging 24 h after cytokine RNA mix injection into naïve mice, (**H**) *ex vivo* measurement of FDG uptake. Significance was determined using a two-tailed t-test for unpaired samples.

Next, we tested cytokine expression of a combination of three formulated RNAs encoding IFN-α, IL-7-Alb and Alb-IL-2mut that has lower affinity to CD25 and therefore decreased capacity to activate T_reg_ (Supplementary Fig. S1E), from here on referred to as ‘cytokine RNA mix’. To assess cytokine expression *in vivo*, the cytokine RNA mix was administered i.v., and cytokine levels were determined in explanted lung and spleen. Highest lung cytokine levels were detected 3–6 hours post-injection (Fig. 1E) with substantially increased cytokine levels in the lung compared to spleen (Fig.1F).

Having demonstrated the feasibility of targeted delivery of RNA-encoded cytokines to the lung, we analyzed their ability to stimulate the immune system using [18F]-Fluoro-2-deoxy-2-D-glucose (FDG) uptake as previously shown for spleen-targeted RNA-LPX vaccination[12]. An increase in FDG uptake was detected 24 hours after cytokine RNA mix injection in lung and lung-draining mediastinal lymph nodes, but not in non-draining popliteal lymph nodes (Fig. 1G, H), indicating an ongoing immune stimulation. Since recombinant IL-2 is known to induce vascular leak syndrome accompanied by inflammatory perivascular infiltrations [13,14], we histologically confirmed that the cytokine mix containing a CD25-low-binding variant of IL-2 did not induce perivascular infiltrations in the lung (Supplementary Fig. S2A). Furthermore, no signs of lung tissue damage, such as widespread lung inflammation, lung edema, hyaline membranes in lung alveolar spaces, or thrombosis in pulmonary vessels as described by other cationic particles [15,16] were detectable (Supplementary Fig. S2A). This is additionally substantiated by bronchoalveolar lavage fluid (BALF) analysis showing no increase of TNF-α (Supplementary Fig. S2B) and neutrophil content (Supplementary Fig. S2C), further potential indicators for particle-induced pulmonary toxicity.

### Cytokine RNA mix induces anti-tumor activity and immune cell response in an experimental B16F10 metastases model

To evaluate anti-tumor activity of the cytokine RNA mix in an experimental metastasis model, we injected C57BL/6 mice i.v. with B16F10-*luc* tumor cells, leading to tumor growth predominantly in the lung. Cytokine RNA mix treatment compared with irrelevant (control) RNA, led to a significant increase in survival (median survival time, MST) and to a reduction of lung tumor burden as shown by BLI (Fig. 2A). To characterize the cytokine-induced changes in the TIME of B16F10-*luc* metastases-bearing lungs, mice were sacrificed 1 day after the third RNA-LPX infusion for flow cytometric analyses (Fig. 2B). Compared with irrelevant RNA, treatment with cytokine RNA mix resulted in increased numbers, proliferation (shown by Ki67 staining) of CD8^+^ T and NK cells and increased activation (CD69 and 4-1BB) of CD8^+^ T cells as shown by flow cytometric analysis (Fig. 2B, C). In an *ex vivo* stimulation assay, CD8^+^ T cells from the lungs of cytokine RNA mix-treated mice showed augmented IFN-γ, granzyme B (GrzB), and IL-2 release and degranulation (CD107a), compared with irrelevant RNA-treated mice (Fig. 2D, E). Boolean gating furthermore demonstrates higher occurrence of polyfunctional CD8^+^ T cells in cytokine RNA mix-treated mice (Fig. 2F). NK cells from lung showed increased GrzB production (Supplementary Fig. S3A) and immunostimulatory effects were also detected in the spleen (Supplementary Fig. S3B, C).

**Figure 2:**
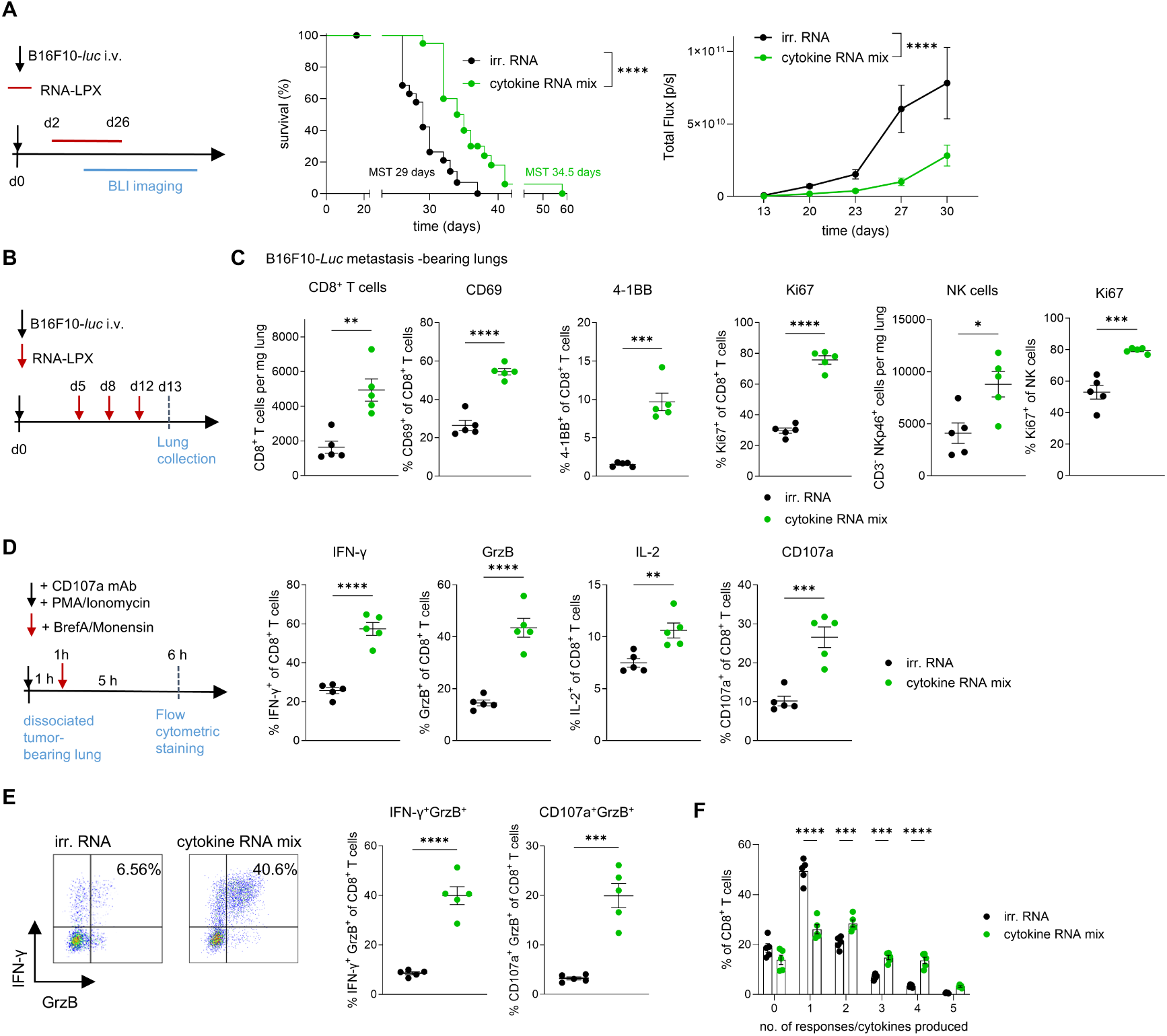
Anti-tumor efficacy and increased activation and effector function of T cells of lung-targeted cytokine RNA mix in B16F10-*luc* metastasis-bearing lungs. (**A**) Experimental design: C57BL/6 (n=20 mice per group) were injected i.v. with B16F10-*luc* cells; subsequently, cytokine RNA mix or irrelevant RNA were administered i.v. as RNA-LPX twice per week for a total of 8 injections from d2 to d26. Mice were monitored until termination criteria were reached. Significance for survival was determined via Mantel-Cox logrank test. BLI was performed at the indicated time points and total flux was quantified in the area of the lung. Significance for total flux was calculated by mixed effects analysis and Geisser-Greenhouse correction (**B**) Experimental design: For flow cytometric analysis, n=5 C57BL/6 mice per group were injected with B16F10-*luc* cells, treated with RNA-LPX on d5, d8 and d12 post tumor injection and sacrificed at d13; quantification is shown in (**C**). (**D**, **E**) Dissociated lungs of treated mice were stimulated with phorbol 12-myristate-13-acetate (PMA)/Ionomycin to analyze degranulation and cytokine release. (**E**) Exemplary IFN-γ and GrzB release by CD8^+^ T cells are shown. (**C**–**E**) Significance for pairwise comparisons was determined by unpaired two-tailed t-test. (**F**) Boolean analysis quantifying the number of CD8^+^ T cells per group with none, one and up to 5 effector molecule responses (CD107a, GrzB, IL-2, TNF-α, and IFN-γ). Significance was calculated based on two-way ANOVA and post-hoc Sidak test for multiple comparisons.

### Orchestration of innate and adaptive immune responses in an experimental CT26 metastasis model leads to long-term survivors

As additional model, BALB/c mice were injected i.v. with CT26 tumor cells followed by RNA-LPX treatment (Fig. 3A). In this model, not only survival was significantly prolonged, but also long-term survivors were obtained (Fig. 3B). To assess long-term immunity, these mice were re-challenged with CT26 tumor cells 98 days after initial tumor induction. While treatment-naïve control mice had to be sacrificed before d20 post tumor induction (p.t.i.), pre-treated long-term survivors did not meet the experimental endpoint criteria until d41 after re-challenge, indicating that an immunological memory preventing tumor cell engraftment had been generated (Fig. 3C).

**Figure 3:**
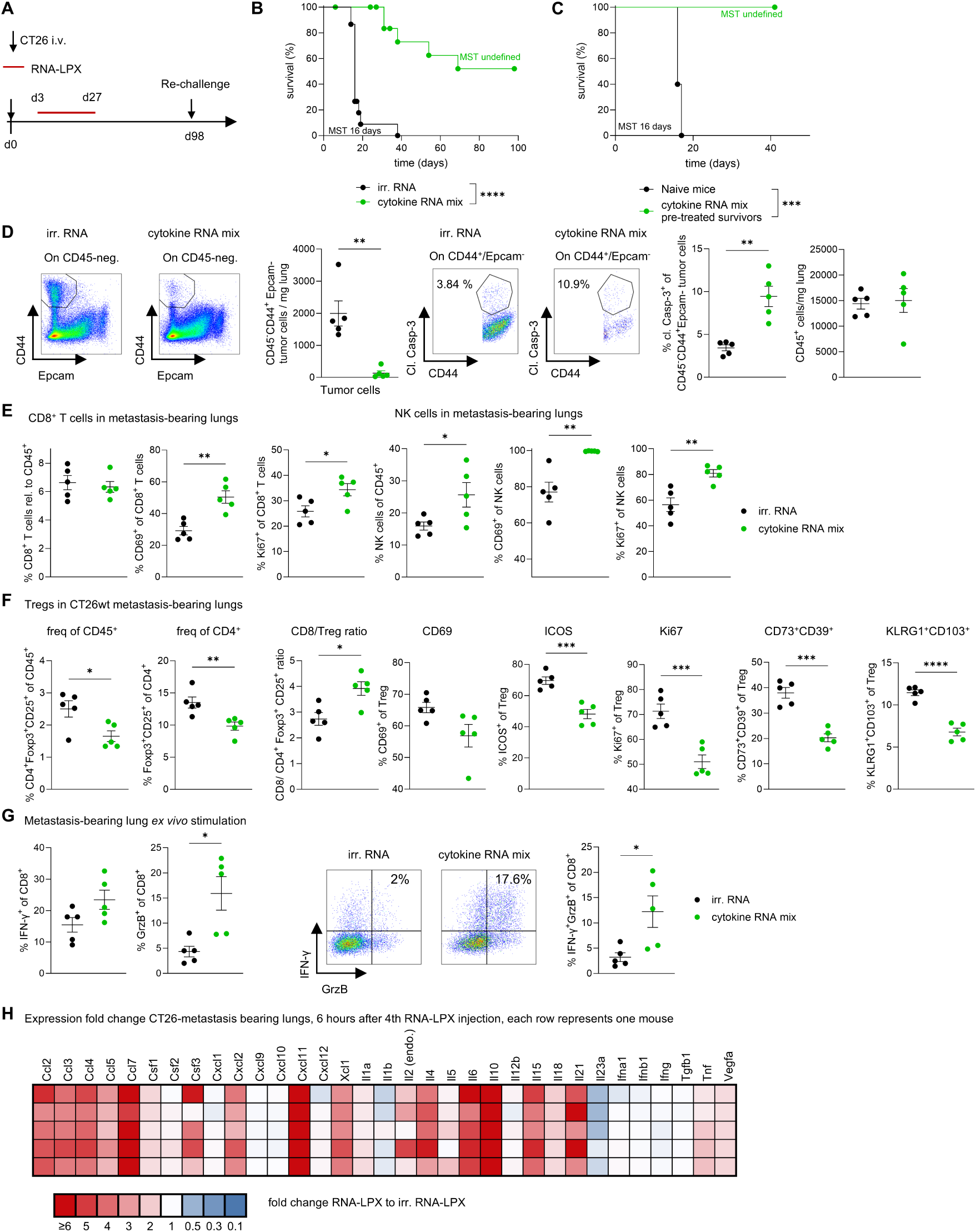
Cytokine RNA mix prolongs survival in a CT26 lung metastasis model accompanied by increased activation of CD8^+^ T and NK cells, reduced T_reg_ and increased effector function. (**A**) Experimental design: BALB/c mice (n=15 per group) were injected i.v. with CT26 tumor cells; subsequently, cytokine RNA mix or irrelevant RNA was administered i.v. as RNA-LPX twice per week for a total of 8 injections from d3 to d27. Survivor mice from the cytokine RNA mix-treated group (n=5) compared to naïve BALB/c mice (n=10) were re-challenged with CT26 tumor cells. (**B**) Survival after the initial i.v. CT26 tumor cell inoculation. (**C**) Survival after re-challenge. Significance was determined via Mantel-Cox logrank. (**D-F**) BALB/c mice (n=5 per group) were injected i.v. with CT26 tumor cells, RNA-LPX was administered i.v. at d3, d6 and d10 p.t.i., mice were sacrificed at d11, lungs were collected and analyzed via flow cytometry. Significance for pairwise comparisons was determined by unpaired two-tailed t-test. Flow cytometric analysis of (**D**) tumor burden, (**E**) CD8^+^ T and NK cells and (**F**) CD4^+^ Foxp3^+^ CD25^+^ T_reg_. (**G**) Functional analysis: after tumor cell inoculation, mice were treated with RNA-LPX at d3, d6, d10 and d13 p.t.i., mice were sacrificed at d14 for an *ex vivo* stimulation assay with PMA/Ionomycin (**H**) qRT-PCR was done to determine fold-change expression of cytokine RNA mix-treated normalized to irrelevant RNA LPX-treated lung samples (n=5 per group indicated in rows) for the indicated cytokines and chemokines.

Flow cytometric analysis of CT26-metastasis-bearing lungs showed a lower burden of tumor cells and higher levels of cleaved caspase-3 in tumor cells indicating apoptotic pathway induction in mice treated with cytokine RNA mix versus control RNA (Fig. 3D). CD8^+^ T cells showed increased activation and increased Ki67 expression (Fig. 3E), although no increase in the relative CD8^+^ T cell fraction among total CD45^+^ leukocytes was observed at this time point. Macrophage and DC subsets, including cross-presenting cDC1, showed an increased expression of co-stimulatory CD86 (Supplementary Fig. S4A) and CD69 (Supplementary Fig. S4B). Compared with irrelevant RNA, cytokine RNA mix treatment resulted in an increase of lung-infiltrating NK cells exhibiting increased activation (Fig. 3E) as well as increased abundance, proliferation and expression of co-stimulatory CD86 on ILC2 and ILC3 in tumor-bearing lungs (Supplementary Fig. S4C).

T_reg_, identified as CD4^+^CD25^+^ FoxP3^+^ cells, were reduced (Fig. 3F), accompanied by a reduction in ICOS and Ki67 expression on T_reg_ and an increase in CD8^+^/T_reg_ ratio. Furthermore, compared with irrelevant RNA, cytokine RNA mix treatment led to a significant reduction of KLRG1^+^ CD103^+^ T_reg_ as well as T_reg_ subsets co-expressing CD39 and CD73 (Fig. 3F), two ectoenzymes that mediate immunosuppressive effects by catalyzing adenosine generation. Because lungs harbor a sizeable CD4^+^Foxp3^+^CD25^−^ T_reg_ population, we confirmed these results by measuring the effects in lung also on CD4^+^Foxp3^+^ T cells (CD25^+/−^; Supplementary Fig. S4D).

*Ex vivo* stimulation of CD8^+^ cells from the CT26 metastasis-bearing lungs of mice treated with cytokine RNA mix showed increased GrzB production and increased IFN-γ^+^ GrzB^+^ double-positive CD8^+^ T cells compared with controls (Fig. 3G). Similar effects can also be observed in spleen (Supplementary Fig. S4E, F).

To conduct a detailed analysis of the cytokines released after cytokine RNA mix treatment, we performed qRT-PCR-based expression analysis of tumor-bearing lungs (Fig. 3H). Lung-targeted cytokine RNA mix treatment led to a marked induction of CXCL11, as well as CCL5 and XCL-1. IL-15 and IL-21 were also upregulated in lungs treated with cytokine RNA mix compared with the control treatment. In conclusion, these findings showcase the multifaceted impact of the cytokine RNA mix treatment on both adaptive and innate immune responses.

### scRNAseq data confirms immunological effects of cytokine RNA mix treatment

To confirm and extend flow cytometric observations, single-cell RNA sequencing (scRNAseq) was performed on CD45^+^ cells isolated from CT26 metastasis-bearing lungs 1 day after the third cytokine RNA mix or irrelevant RNA treatment, and 22 clusters were defined (Fig. 4A, Supplementary Fig. S5). Among the three CD8^+^ T cell clusters, CD8^+^ T cells cluster 1 (CD8-1) had the highest *Il7r* expression and a naïve/memory phenotype based on expression of *Ccr7*, *Tcf7*, *Lef1, Sell,* and no expression of effector cell–related genes, such as *Cd44*, *Fas*, *Cxcr3* (Supplementary Fig. S6A). Clusters CD8-2 and CD8-3 showed low *Sell* expression and expression of *Cd44* and *Cxcr3* indicating an effector phenotype (Supplementary Fig. S6A). The CD8-1 and CD8-2 clusters were reduced (Fig. 4B), while the effector-like CD8-3 cluster shows much higher frequency upon treatment with cytokine RNA mix compared with irrelevant control RNA (17-fold, Supplementary Fig. S6B, C) and expressed the highest level of *Cd27*, *Itgal*, *Cxcr3, Cd69* and *Gzmb* (Supplementary Fig.6A). NK cell subsets were separated into 4 clusters (NK1-3, NK-like, Supplementary Fig. S6A, D). NK2 (*Klrk1*/NKG2D^high^ IL2rb^high^) and NK3 (*Top2a*^high^ *Mki67*^high^ proliferating NK) were enriched upon treatment with cytokine RNA mix, whereas the NK1 cluster (*Klrc1*/NKG2A^high^) decreased in frequency (Fig. 4B, Supplementary Fig. S6C). NK2 harbored the highest expression of *Itgam*, *CD69*, *IL2rb*, *Ifnar1*, *Sell* and *Spn* (Supplementary Fig. S6A, D) indicating an activated mature phenotype. The proliferating NK3 cells showed a remarkable increase of genes related to glycolysis, citric acid cycle (TCA) and electron transport chain (ETC) compared with clusters NK1 and NK2 (Supplementary Fig. S6E). This increase is presumably linked to meeting the high energy demands during proliferation

**Figure 4:**
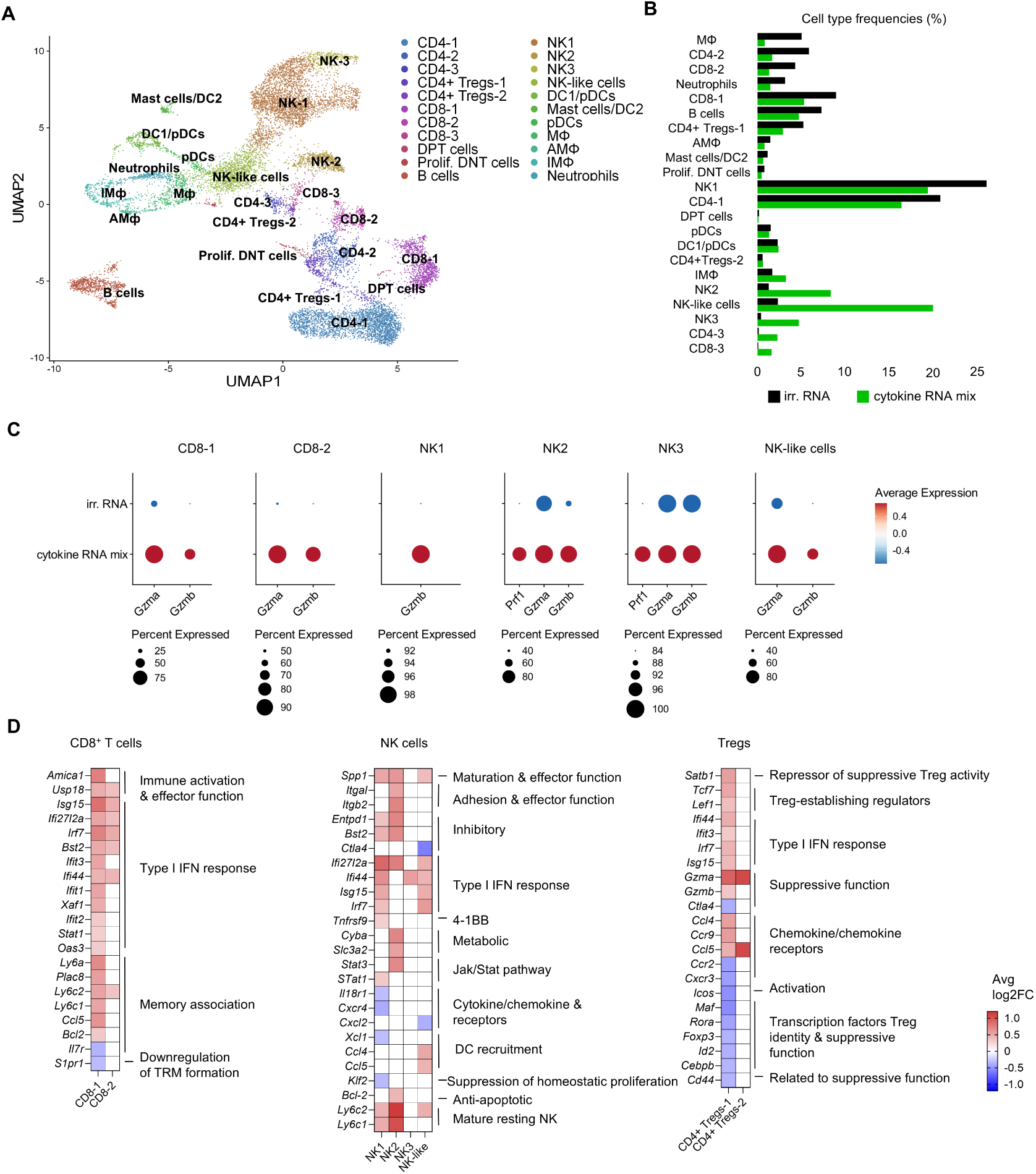
scRNAseq analysis confirms that cytokine RNA mix treatment results in the activation of CD8^+^ T cells and NK cells, and has a suppressive effect on T_reg_. Experimental design: BALB/c mice (n=5 per group) were injected i.v. with CT26 tumor cells; cytokine RNA mix or irrelevant RNA was administered i.v. at d3, d6 and d10 post tumor cell injection, mice were sacrificed at d11, lungs were collected and pooled at same ratios before sorting of CD45^+^ cells. CD45^+^ cells were subjected to scRNAseq. (**A,B**) Uniform manifold approximation and projection (UMAP) of 22 assigned clusters (**A**) and cell type frequencies (**B**) of CD45^+^ cells isolated from the lungs of cytokine RNA mix-treated versus irrelevant RNA-treated mice. (**C**) Effector function visualized as bubble plot. (**D**) Selected differentially expressed genes in cytokine RNA mix-treated versus irrelevant RNA-treated samples are shown as average log2 fold change. Note only significantly expressed values (p ≤ 0.05) are shown in colouring (blue = downregulated, red = upregulated), non-significant values are set to “0”/white.

In line with flow cytometric observations, increased expression of effector molecules was detected for both CD8^+^ T cell (CD8-1, CD8-2) and all NK cell subsets (Fig. 4C). Overall, NK cells were identified as major source of *Gzmb* and *Prf1* in the TIME (Supplementary Fig. S7A).

Next, we aimed to understand the role of type I and type II interferons in the mode of action of the cytokine RNA mix. In contrast to *Gzmb*, *Ifng* was only modestly upregulated after treatment with cytokine RNA mix (Supplementary Fig. S7B) and did not reach significance in CD8^+^ T or NK cell differentially expressed gene analysis, which is in line with the flow cytometric and qRT-PCR results (Fig. 3G, H). Notably, a neutralization experiment confirmed that IFN-γ does not appear to be a major driver of the anti-tumor efficacy (Supplementary Fig. S7C, D). Analysis of differentially expressed genes (Fig. 4D, Supplementary Fig. S8) showed upregulation of type I interferon response genes (for example *Ifi44*, *Ifit1*, *Ifit2, ISG15*) on all NK subsets, CD8-1-2 and T_reg_ as expected when delivering IFN-α to the TIME. Type I interferon has been described to boost both NK and T cell–mediated effector functions, while suppressing T_reg_[5]. This is in line with increased expression of markers associated with effector function and maturation in NK and CD8^+^ T cell subsets (Fig. 4D). Furthermore, factors associated with increased suppressive function (e.g. *Ctla4*) and T_reg_ identity (e.g. *Foxp3*, *Maf*, *Rora*) were downregulated in T_reg_ (Fig. 4D), whereas other inhibitory factors (e.g. *IL10*, *Lag3*, *Pdcd1*, *Tgfb1*, *Il4)* were not significantly regulated on T_reg_. Despite a strong type I IFN response, IFN-α–encoding RNA-LPX alone had no relevant anti-tumor efficacy, nor had Alb-IL-2mut and IL-7-Alb alone but showed clear synergy for survival as well as in modulation of the TIME with IL-7-Alb and Alb-IL-2mut in the CT26 (Supplementary Fig. S9A-C) and B16F10 i.v. tumor model (Supplementary Fig. S9D-E). Comparison of the immunological effects of the full cytokine mix therapy to Alb-IL-2mut and IL-7-Alb alone in flow cytometric analyses highlighted the role of IFN-α on Treg suppression as well as on CD8^+^ T cell functionality (Supplementary Fig. S9).

Overall, no marked induction of co-inhibitory receptors opposing effector function was observed in the scRNAseq analysis (Supplementary Fig. S10A), flow cytometry or immunofluorescence staining (Supplementary Fig. S10B-E).

In summary, the scRNAseq analysis further confirms the enhanced effector functions of CD8^+^ T cells and NK cells, while showing a reduction in T_reg_-mediated suppression. Additionally, the data suggest regulation of genes associated with T cell memory and underscores the significance of type I interferon in the cytokine RNA mix.

### CD8^+^ T and NK cells mediate the anti-tumor effect of the cytokine RNA mix, leading to efficacy in mouse models of T cell escape

Having shown increased activation and effector function of CD8^+^ T and NK cells, we investigated whether both cell types are crucial mediators of the therapy-induced anti-tumor response. *In vivo* depletion of CD8^+^ T cells or NK cells prior to treatment led to significantly reduced survival (Fig. 5A–C). Logrank test with an emphasis on early or late differences showed that the absence of CD8^+^ T cells had the largest impact on survival in the late phase (Fig. 5B), whereas the absence of NK cells was most relevant in the early phase (Fig. 5C). Immune escape is a common event and poses a major challenge in cancer therapy. Given the dual activation of innate and adaptive immunity that we observed, we hypothesized that the cytokine RNA mix treatment may also be efficient in models of T cell escape. Using a CT26 knockout cell line either devoid of the immunodominant T cell antigen gp70 (CT26gp70 k.o.; model for antigen loss) or β-2 microglobulin (B2M) (CT26B2M k.o.; model for MHC-I loss) we confirmed a substantial survival benefit of treatment with cytokine RNA mix in both models (Fig. 5D). Furthermore, upon cell depletion in the B2M k.o. tumor model, NK cells appeared to mediate the anti-tumor effect (Fig. 5E) with a crucial role of the activating receptor NKG2D (Fig. 5E). In summary, these findings indicate that the effectiveness of the lung-targeted cytokine RNA mix treatment is not confined to the presence of a specific T cell antigen but can also be achieved in an effector T cell-independent manner through NK cells, confirming its efficacy in models of T cell escape.

**Figure 5:**
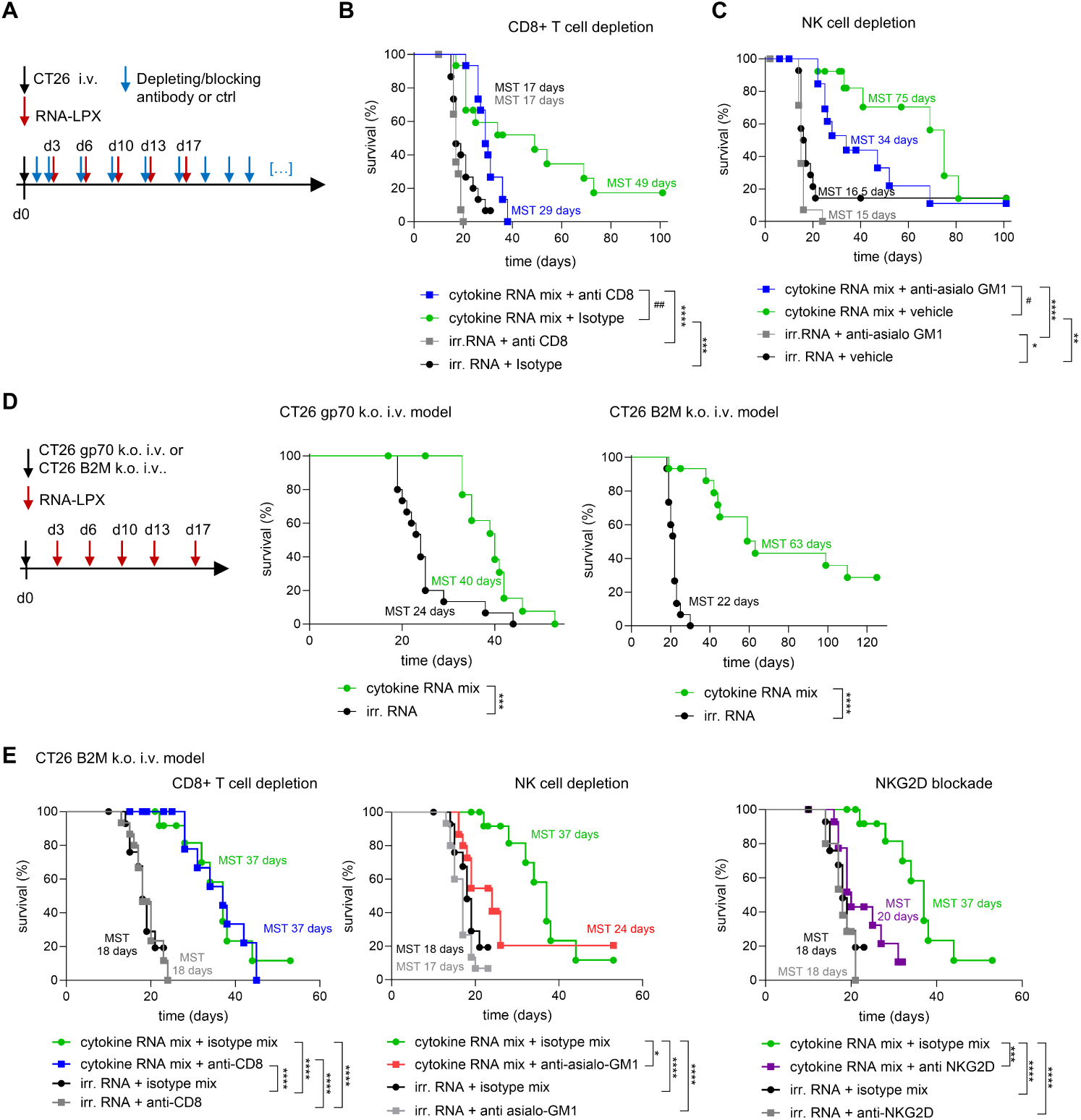
The anti-tumor effect of cytokine RNA mix is mediated by CD8^+^ T cells and NK cells and is maintained in models of tumor T cell escape. (**A**) Experimental design for (**B-C;** CT26 tumor model) and (**E;** CT26 B2M k.o. tumor model); n=15 BALB/c mice per group. Depletion or blocking antibody treatment was started 2 days prior to RNA-LPX treatment to ensure depletion before treatment start. (**B, C**) Survival according to termination criteria. (**D**) Experimental design and survival in CT26B2M k.o. or CT26gp70 k.o. tumor cells i.v. tumor model, n=15 mice per group. (**E**) BALB/c mice injected i.v. with CT26B2M k.o and depletion/neutralization antibodies were applied as shown in (**A**). Note that data in (**E**) were generated within the same experiment, irrelevant RNA + isotype mix as well as cytokine mix RNA + isotype mix refers to the same groups in all 3 plots. Survival was analyzed via Mantel-Cox logrank test. For (**B**) and (**C**), groups 1 and 2 were furthermore compared via logrank test with emphasis on early and late differences, (**B**) ## Logrank test with emphasis on late differences ((rho=0, lambda=1): group 1 vs 2: p=0,00235; (**C**) # Logrank test with emphasis on early differences (rho=1, lambda=0): group 1 vs 2: p=0,0378.

Next, we explored whether combining the cytokine RNA mix with cisplatin and pemetrexed would result in a synergistic effect and observed a strong tendency towards synergy along with increased activation of NK and CD8^+^ T cells (Supplementary Fig. S10F-H).

### Primary patient-derived tumor culture indicates translatability of the cytokine RNA mix

To evaluate the effect of cytokine RNA mix treatment in an *ex vivo* model of human cancer, we cultured tumor punches derived from patients with NSCLC for 48 hours with supernatants of HEK cells lipofected with cytokine RNA mix or with irrelevant RNA (Fig. 6A) and assessed treatment response of the tumor-infiltrating lymphocytes. Flow cytometric analysis revealed a reduction in CD45^−^ cells (Fig. 6B) and increased abundance and activation of CD8^+^ T cells, as well as an increase of tissue-resident memory T cells (Fig. 6C). Increased abundance of CD8^+^ T cells and increased numbers of GrzB–expressing CD8^+^ T cells was confirmed by immunofluorescence (Fig. 6D, E). Consistently, increased levels of GrzB could be detected in supernatants upon treatment with cytokine RNA mix (Fig. 6F). These results indicate that cytokine RNA mix was able to trigger immune stimulatory effects in the human TIME in a patient-derived 3D *ex vivo* model, suggesting a translational relevance for lung malignancies.

**Figure 6:**
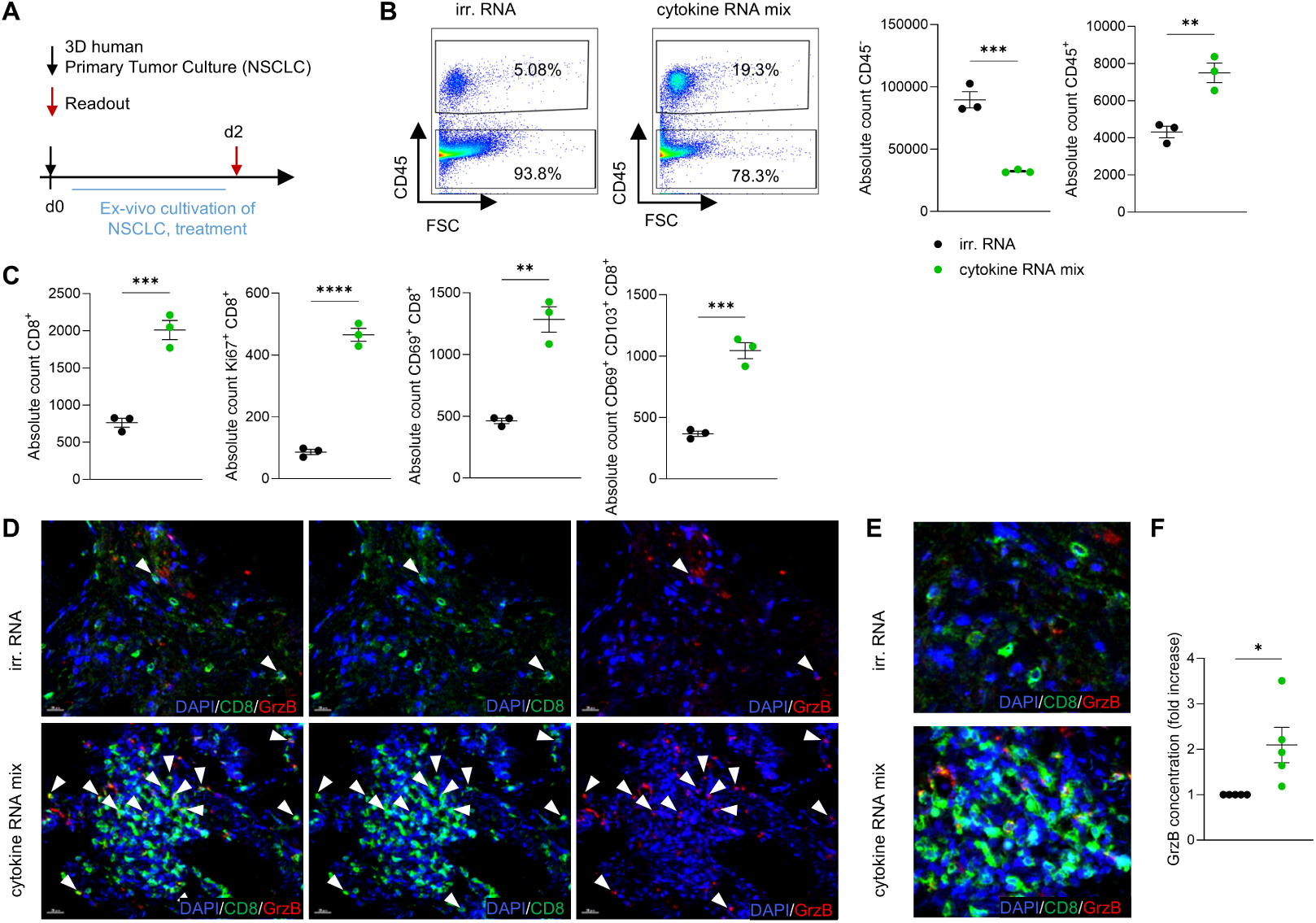
Patient-derived NSCLC ex vivo culture shows cytokine mix-mediated CD8^+^ T cell activation and GrzB release. (**A**) Experimental design: NSCLC was resected, cultured *ex vivo* for 48 hours with supernatants of HEK cells lipofected with cytokine RNA mix or irrelevant RNA. (**B**) Flow cytometric quantification of CD45^+^ and CD45^−^ cells. (**C**) Quantification of CD8^+^ T activation (CD69, CD25) and tissue resident memory T cells (CD69^+^ CD103^+^). (**B, C**) One donor with technical replicates is shown, similar results were observed in n=3 independent donors. (**D, E**) Multiplex immunohistochemistry showing nuclei (blue), CD8 (green) and GrzB (red). Arrows indicate co-localization of CD8 and GrzB. (**F**) Fold increase (cytokine-treated/irrelevant) of GrzB secretion as determined in supernatants (similar results were observed in 2 independent donors). Significance for pairwise comparisons was determined by unpaired two-tailed t-test.

## DISCUSSION

In cancer immunotherapy, local delivery is a strategy used to obtain high concentration of therapeutic agents at the tumor site. Earlier attempts included intratumoral cytokine injection, an approach that has been further advanced by the use of mRNA-encoded cytokines [17]. A major limitation of this strategy is the need for tumor to be accessible, which is often challenging for lung metastases or primary lung tumors. Albumin fusions or conjugates of proteins have been shown to increase circulatory half-life via reduced renal clearance and induce accumulation at the sites of vascular leakiness or regions rich with proliferating tumor cells, also described as enhanced permeability and retention effect [18]. Our prior work showed that modified RNA encoding albumin-fused cytokines formulated to target the liver, increased cytokine persistence thereby promoting strong antitumoral therapeutic effects [7,19]. In line, we demonstrated in this study the feasibility of delivering albumin-fused cytokines to the lung using the previously described cationic LPX [8–10]. Notably, we used a CD25 binding-deficient IL-2 variant to avoid endothelial stimulation via CD25 [14] and thus mitigate the risk of vascular leak syndrome and pulmonary edema as well as reducing stimulation of T_reg_.

In the B16F10, known as a poorly immunogenic tumor model [20], treatment with cytokine RNA mix resulted in a significant increase in CD8^+^ T cell numbers compared with irrelevant RNA treatment. We did not observe the same effect in the CT26 metastasis model, which is highly immunogenic [20]; however, in this model, scRNAseq analysis identified NK cell subsets as major producers of *Gzmb* in the lung TIME.

Complementary qRT-PCR profiling showed that recruitment of effector cells may have been facilitated by an inflammatory response and the release of chemokines, namely CXCL11, known for its role in CD8^+^ T cell activation and recruitment [21], XCL1 and CCL5 that are capable of recruiting cross-presenting cDC1[22], as well as IL-15 and IL-21 that contribute to the activation, expansion and effector function of CD8^+^ T and NK cells [23,24].

Surprisingly, IFN-γ did not play a major role in driving the anti-tumor response, as demonstrated by a neutralization experiment. IFN-γ is known to upregulate MHC-I [25], which can improve response to CD8^+^ T cells, whereas it decreases sensitivity for NK-cell mediated lysis. Since both CD8^+^ T and NK cells were involved in the response to cytokine RNA mix treatment, the release of IFN-γ in this specific setup might not have clear pro- or anti-tumor effects consistent with published pro- and anti-tumor effects of IFN-γ [26].

Type I IFN, instead, appeared to be crucial for treatment efficacy. Macrophage and DC subsets showed increased activation and expression of co-stimulatory CD86 with cytokine RNA mix treatment versus irrelevant treatment, potentially supporting increased T cell priming and especially cross-priming – which is consistent with previous reports on activities of type I IFN [27]. Additionally, type I IFN is known to inhibit T_reg_ and attenuate their suppressor function [28]. Consistently, an improved CD8^+^/T_reg_ ratio in CT26-metastasis-bearing lungs was observed via flow cytometry and substantiated by further flow cytometric analyses, as well as scRNAseq data.

The activity of type I IFN on CD8^+^ T cells has been described as a crucial mediator of effector and memory responses in viral infections [29] and can sensitize T cells to IL-2–mediated proliferation [30] fitting our observation of increased Ki67 expression on CD8^+^ T cells and an improved CD8/T_reg_ ratio comparing the full cytokine RNA mix to Alb-IL-2mut/IL-7Alb in the CT26 tumor model. Most importantly, we demonstrated a clear synergy of the cytokines for therapeutic efficacy. IL-7 plays a pivotal role in regulating memory T cell homeostasis [4]. Despite the relatively early time point of the scRNAseq analysis, at which memory formation is unlikely, genes associated with memory (e.g. *Ly6c, Plac8)*[31,32], longer life span (*Bcl-2*) [33], increased functionality upon re-stimulation, and tissue-resident memory (TRM) formation [34] (downregulation of *S1pr1*), appeared to be upregulated in CD8-1. Immune protection observed in the re-challenge experiments indicated that indeed a long-term memory response was successfully established.

Treatment of human T cells with type I IFN has been shown to increase the expression of co-inhibitory receptors [35], in line with other reports indicating that, during a chronic viral infection, prolonged IFN-I signaling may promote T cell exhaustion [36]. Despite IFN-α in our cytokine RNA mix, we did not observe an upregulation of co-inhibitory receptors on CD8^+^ T cells via flow cytometry or by scRNAseq analysis. The reason may be that the delivered IFN-α is expressed temporarily at a low dose that does not reach the level of chronic exposure linked to the functional or metabolic exhaustion of CD8^+^ T cells [37]. Type I interferons are also known to have a positive impact on NK cell function, homeostasis and maturation via direct or indirect mechanisms [38]. IFN-α can stimulate DCs to express IL-15 but also has direct effects on NK cells. Furthermore, IL-2 can drive NK cell expansion [39], which may explain our observations in the flow cytometry and scRNAseq analyses.

Tumor T cell escape is a recurrent event caused by the loss of MHC-I [40] or relevant T cell antigens [41], which may present a major risk of therapy failure. In our study we addressed both aspects of T cell escape by utilizing experimental metastases models employing transgenic CT26 tumor cells that lacked the expression of either MHC-I (B2M k.o.), or the immunodominant antigen (gp70 k.o.). In both models, we observed a significant survival benefit that could be attributed to NK cells, with a predominant role of the NKG2D axis in the CT26 B2M k.o. model. Anti-tumor activity of CD8^+^ T cells in an NKG2D-dependent manner has been described for MHC-I deficient tumors[42]. In this study, we identified CD8-2 and CD8-3 as NKG2D (*Klrk1*) positive cluster (Fig. 5c). However, anti-tumor efficacy in the B2M k.o. tumor model was not abrogated in the absence of CD8^+^ T cells, suggesting that NKG2D-positive NK cells were the crucial effector cells mediating lysis of MHC-I-deficient tumor cells.

We also tested the immune-stimulatory effects of the cytokine RNA mix in an *ex vivo* culture of primary human tumor fragments. Despite the lack of circulation and therefore lack of supply of new leukocytes, but preserving the endogenous resident leukocytes, we confirmed the cytokine RNA mix-mediated T cell activation and observed signs of proliferation of tumor-infiltrating CD8^+^ T cells; the immune activation was further substantiated by increased GrzB release.

Overall, our data indicate that, in mouse experimental metastases models, lung-targeted delivery of an RNA-encoded mix of IFN-α, IL-7-Alb and Alb-IL-2mut led to an early activation and proliferation of NK cells, which were a main source of *Gzmb* in the lung TIME and, together with CD8^+^ T cells, contributed to an anti-tumor immune response. The response was facilitated by T_reg_ inhibition in the lung as well as by myeloid activation that may in turn have led to increased T cell priming. Interestingly, cytokine RNA mix treatment activated CD8^+^ T cells, which are mainly relevant for the later phase of survival, but appeared to be dispensable in a MHC-I k.o. model, where the anti-tumor response was mediated by NK cells. Thus, we provide evidence for an i.v. administered, locally-active cytokine mix that engages both innate and adaptive immunity to boost the endogenous immune response, thereby leading to anti-tumor efficacy in MHC-I-proficient and deficient tumor models. Our study presents a comprehensive analysis on the immunologic mode of action of the RNA-encoded cytokine combination supporting the clinical evaluation of the lung-targeted RNA-encoded cytokine mix.

## METHODS

### Ethic statement

All human tissue samples were provided by the tissue bank of the University Medical Center Mainz in accordance with the regulations of the tissue biobank and the approval of the ethics committee of University Medical Center Mainz.

### RNA

The following RNAs were used (h: human; m: murine): hAlb-hIL-2mut, hIL-7-hAlb and hIFN-a2a (Fig. 1E,F, S1E, Fig. 6/*ex vivo* experiments using human (tumor) samples) or hAlb-hIL-2mut, mIFN-a4 and mIL-7-mAlb for (therapeutic) experiments performed in mice. As irrelevant control RNA, a mix of empty vector RNA (involving backbone elements as a secretion signal and MHC-I trafficking domain as published in [43]) and RNA-encoded Alb was used at an RNA mass ratio of 1:2 empty vector to Alb-encoding RNA. The cytokines or cytokine-albumine fusions of interest were cloned into plasmid vectors containing sequences for the 5’ untranslated region (UTR), a 3’ UTR and 100 nt poly A tail interrupted by a 10 nt linker (A30LA70) [44–46]. RNA was generated from a linearized DNA template by *in vitro* transcription (IVT) with T7 RNA polymerase (Thermo Fisher Scientific) in the presence of adenosine 5′-triphosphate, cytidine 5′-triphosphate, guanosine 5′-triphosphate and 1-methylpseudouridine-5’-triphosphate (m1ΨTP; Thermo Fisher Scientific) substituting for uridine-5’-triphosphate (UTP) as previously described [11,47]. Depending on the construct a cap was either introduced co-transcriptionally (CleanCap N-7413, TriLink Biotechnologies) or after IVT by enzymatic capping (NEB vaccinia capping system; eCap1). To remove double-stranded RNA contaminants, cellulose purification was performed as previously described [47]. For details on RNA-LPX, see supplemental information.

### RNA-LPX

Solutions of L5 Liposomes (1:1 DOTMA: Cholesterol) were diluted in water to the appropriate concentration (to achieve a molar N/P ratio of 4:1 cationic lipid to RNA in the mixed LPX solution) and vortexed by hand. Simultaneously, RNA was diluted in NaCl and vortexed. These were kept on ice until needed. These 2 phases were loaded into syringes and connected to a fluid path system to ensure defined mixing conditions. The first fraction of the solution were discarded before collection of the LPX was started. In a second step, the LPX were mixed with a NaOAc and Sucrose-containing storage matrix and stored at 2-8°C until needed. Where applicable, RNA-LPXs were gently vortexed by hand to achieve the appropriate ratio and concentration. QC was performed using dynamic light scattering (DLS) showing a typical size of 150-250 nm.

Exceptions are Figures 1 C, D and Suppl. Fig. 1C, D where RNA-LPX containing the same lipid composition and lipid: RNA ratios as above were manufactured as follows:

Fig. 1C, D: As previously described (*21-23*), the RNA-lipoplexes were generated by manual dilution of RNA in H2O and NaCl prior to addition of liposomes to obtain a final RNA concentration of 0,05 mg/ml.

Suppl. Supplementary Fig. S1 C, D: RNA was added to water, followed by NaCl and vortexed. Liposomes were then added with a pipette to achieve an RNA-LPX in NaCl, at the appropriate concentration (a molar N/P ratio of 4:1 cationic lipid to RNA in the mixed LPX solution). The RNA-LPX was then diluted to an RNA concentration of 0.1 mg/mL and stored at 2-8°C until needed. Particles were measured using DLS, as to be typically 150-400 nm.

Zeta potential for RNA-LPX was typically 40 mV +/− 10 mV.

### Quantification of cytokine levels in the lung and spleen by ECLIA assay (hAlb hIL2mut and hIL7-hAlb) or ELISA assay (hIFN-α2a)

Whole lung and spleen were collected 3, 6, 24, and 72 hours after the injection and transferred to a pre-weighed and labeled homogenizing tube (CK Mix, BertinCorp, Rockville (MD), USA). Weight of the organs/tissue samples was recorded, and the tubes stored at −80°C. To obtain tissue lysates, samples were processed with Precellys^®^ Evolution (VWR™, Darmstadt, Germany).

Concentrations of translated hAlb-hIL2mut and hIL7-hAlb protein in tissue lysates were determined by electro-chemiluminescence immunoassays (ECLIA) using custom-developed reagents (Meso Scale Discovery). hAlb-hIL2mut and hIL7-hAlb were captured in wells coated with monoclonal anti-human IL-2 or anti-human IL-7 antibodies, respectively. Captured cytokines were detected by using another monoclonal anti-human cytokine antibody conjugated to an electro-chemiluminescent label (SULFO-TAG). Luminescence signal of the label was measured using a MESO QuickPlex SQ120 imager.

hIFN-α2a concentrations in tissue lysates were determined using the VeriKine^TM^ Human IFN Alpha ELISA Kit (PBL assay science; cat. no. 41100).

### PET imaging

BALB/c mice were injected i.v. with RNA-LPX and imaged on subsequent days by FDG-PET imaging. Injection of [^18^F]FDG were performed on anesthetized animals with 2.0% isoflurane (Abbott, Wiesbaden, Germany) evaporated in oxygen at a flow rate of 0.5 L/min. Immediately before the injection of [^18^F]FDG body weights of the mice were measured. In total, ∼5 MBq [^18^F]FDG dissolved in 100 µL 0.9% NaCl solution was injected i.v.. Whole body PET scans were performed after injection of [^18^F]FDG after 45 minutes using a dedicated Focus 120 small-animal PET scanner (Concorde Microsystems/ Siemens Preclinical Solutions, Knoxville, USA), for 12 min while the mice were kept under anesthesia. Reconstruction was performed in microPET Acquisition Workplace V02.3200 with OSEM3D followed by map. The reconstructed voxel size was 0.432×0.432×0.796 mm^3^. The percentage of injected dose per cubic centimeter (%ID/cm³) values were calculated as follows: mean activity in VOI/(injected activity·10^6^)/100.

### [^18^F]FDG biodistribution

In addition to *in vivo* PET data, organs were analyzed by γ-counting *ex vivo* in separate groups, 55 min after tracer injection. A standard solution with a known radioactivity served as reference to calculate the percentage of injected dose per gram of tissue (%ID/g). Tubes containing the standard solution and the organs were measured via Wizard2 automated γ-counter (Perkin Elmer) at an energy window of 350 to 650 keV. The resulting decay corrected counts per minute were then first normalized to the injected dose with the help of the standard solution and to the respective weight of the organ to obtain %ID/g.

### Analysis of STAT5 phosphorylation upon hAlb-hIL2 vs. hAlb-hIL2mut treatment

Peripheral blood mononuclear cells (PBMCs) were isolated from human buffy coats by density gradient centrifugation and rested in Iscove’s Modified Dulbecco’s Medium (IMDM, Life Technologies GmbH, cat. no. 12440-053) supplemented with 5% plasma-derived human serum (PHS, One Lambda Inc., cat. no. A25761) for 1 hour (37°C, 5% CO_2_). Phosphatase inhibitor (PhosSTOP, Roche, cat no. 04906845001) was added to the cells after the incubation to obtain a final concentration of 1x PhosSTOP. PBMCs were seeded in a 96-well V-bottom plate (1.5 × 10^5^ cells/well) and incubated for 15 minutes (37°C, 5% CO_2_) with supernatants of HEK293/T7 cells lipofected with RNA constructs encoding hAlb-hIL2mut or hAlb-hIL2. Fixable viability dye eFluor 780 (Thermo Fisher Scientific, cat. no. 65-0865-18) was added during the last 5 minutes of the incubation, after which cells were fixed in 2% formaldehyde fixative (Carl Roth GmbH, cat. no. P087.4) on ice for 10 minutes. Cells were washed and incubated in 100% methanol for 30 minutes at 2–8°C, washed again, and stained with anti-human CD4 (BD, cat. no. 565997), anti-human CD25 (BD, cat. no. 560503), anti-human FoxP3 (Thermo Fisher, cat. no. 17-4776-42), and anti-STAT5/pY694 (BD, cat. no. 612598) antibodies in 1× permeabilization buffer (Thermo Fisher, cat. no. 00-8333-56) for 30 minutes at 2–8°C. STAT5 phosphorylation in CD4^+^CD25^+^FoxP3^+^ regulatory T cells (T_reg_) was analyzed by flow cytometry.

### Animals

Balb/cJRj and C57BL/6j wildtype female mice were purchased from ENVIGO RMS GmbH, Netherlands and Janvier Laboratories, France, respectively with a typical age of 8 weeks. All experiments were approved by the local authority of Rhineland-Palatinate, Germany for the ethical evaluation of animal experiments and animal welfare. All experiments were performed according to Directive 2010/63/EU, animal care was in accordance with institutional guidelines.

Mice were randomly assigned to active treatment or control groups. Depending on tumor model and setup, typically n=10–20 mice were used per group for survival analysis and n=5 for flow cytometric analysis, as indicated in the figure legends. Group size planning was typically performed by g*power based on previously determined expected biologically relevant effect sizes and approved by the local authority.

Termination criteria for *in vivo* experiments were pre-defined (as e.g. weight loss > 20%, body conditioning score 2, noticeably difficult breathing/increased abdominal breathing, criteria based on general condition and behavior of the mouse as judged via a pre-defined scoring system) and approved by the local animal regulation authority. Survival was determined according to pre-defined termination criteria mentioned above and lung tumor burden was macroscopically assessed to confirm that the mouse met the experimental endpoint. Mice were censored in survival curves when the reason for sacrifice or death was not linked to lung tumor burden. Symbols in survival curves are plotted for events as well as censored data.

### In vivo Bioluminescence imaging (BLI)

For in vivo Bioluminescence imaging of mice injected with luc expressing tumor cell lines or LPX, 150mg/kg luciferin were applied intraperitoneally. 5 min after application BLI was performed in „Xenogen IVIS® Spectrum“(Caliper LifeSciences, Hopkin, MA, USA). Mice were kept under inhalation anesthesia with 2.5 % isoflurane during the imaging. Determination of luminescence signal was performed with IVIS Living Image 4.0 Software (Caliper LifeSciences, Hopkin, MA, USA), by quantification of total flux (photons/sec) in manually defined regions of interest (ROI).

To visualize the intensity of the Luc signal in living mice, radiance (photons/[s cm2 sr]) was displayed as a colour-scaled image.

### Tumor cell lines

Colon Carcinoma cell line CT26 was purchased from ATCC (CT26: CRL-2638, lot no. 58494154, female). CT26 with knockouts of B2M and gp70 were generated as described before [7]. Wildtype CT26 as well as CT26 gp70 knockout cell line were cultured in RPMI 1640 medium (Gibco) with 10% FBS (Bio & Sell). For the culture of CT26 B2M k.o. gRPMI 1640 Glutamax (Gibco) with 10% FBS was used. B16-F10 melanoma cell line was purchased from ATCC (CRL-6475, Lot 63048505, male), Luc expression was engineered as described before (*22*) and kept in RPMI 1640 supplemented with 10% heat inactivated FBS, 1% Sodium Pyruvat (Gibco), 10nM HEPES (Gibco), 1% MEM Non-Essential Amino Acids (Gibco) and 0,5µg/ml Puromycin (InvivoGen). Banked cell lines were confirmed to be negative for mycoplasma (Eurofins) and re-authenticated (Microsynth, DSMZ, CLS), all cell lines were cultivated at 37°C and 5% CO2. After cryopreservation cell lines were passaged 3-8 times before i.v. injection in vivo.

### Tumor models and treatments

For therapeutic tumor models Balb/cJRj mice were injected i.v. into the tail vein with 4 x10^5^ CT26, CT26 gp70 k.o or CT26 B2M k.o. tumor cells. C57BL/6j were injected i.v. with 3 x10^5^ luc expressing B16F10. Treatment starts were depending on the experiments between day 2 and day 6 after tumor injection as indicated in figures and figure legends. RNA-LPX for individual cytokines were mixed in a distinct ratio under sterile and RNAse-free conditions directly before administration. 30µg cytokine RNA mix-encoding or irrelevant RNA-LPX containing a total of 30 µg of RNA at the defined RNA mass ratio of 4:1:4 for hALB-hIL-2mut: mIFN-α4: mIL-7-mAlb relating to a molar ratio of ca. 1.5: 1: 1.5 were injected i.v. twice a week into the tail vein.

Depleting in vivo antibodies and corresponding isotypes were intraperitonally injected twice a week up to 30x beginning at day 1 after tumor inoculation. Per injection 200µg anti CD8 (BioXCell) (loading dose 400µg), 20µl anti asialo-GM1 (Wako), 200 µg anti-NKG2D (BioXCell) (loading dose 400µg) or 200µg corresponding isotypes (BioXCell) were administered. For IFNγ neutralisation, antibody or corresponding isotype was intraperitonally injected three times per week up to 30x beginning at day 1 after tumor inoculation with a loading dose of 400 µg followed by 250 µg of anti-IFNγ antibody (BioXCell).

Animals included in survival analysis were sacrificed when pre-defined criteria (see study design) were reached. To quantify the lung nodules sacrificed mice were injected intratracheally with 20% ink solution and lungs were collected. After de-staining in Fekete solution (100 ml 70% ethanol, 10 ml 37% formaldehyde solution and 5 ml glacial acetic acid) and fixation in 4% Histofix (Carl Roth) metastases were counted manually.

### Collection of Bronchioalveolar Lavage Fluid (BALF)

To exclude toxicity of the LPX particles, Balb/cJRj mice were intravenously injected with 200 µl of cytokine RNA mix RNA-LPX, irrelevant RNA-LPX, PBS or intranasally treated with LPS (O55:B5, Sigma-Aldrich, cat no. L2880-10MG) at a dose of 3 mg/kg under Ketamine (120 mg/kg)/ Xylazin (16 mg/kg) anesthesia. For the collection of BALF, mice were euthanized 6h, 24h, 3 days or 6 days after treatment by overdosing with Ketamine 200 mg/kg and Xylazin 20 mg/kg and a subsequent exsanguination via vena cava. An intratracheal probe was used to flush the lungs three times with 0,5 ml of proteinase inhibitor cocktail (Sigma-Aldrich, cat no.11697498001)-containing PBS. The collected fluid per mouse was pooled and centrifuged at 400 x g, the cell pellet was used for flow cytometry and the supernatant frozen for cytokine ELISA.

### Cytokine and total protein quantification from BALF

Cytokines in broncheoalveolar fluid were measured by TNF-α ELISA (Invitrogen, cat no. 88-7324-88) according to the manufacturer’s instructions.

### Tissue preparation

Spleens and lungs from Balb/cJRj and C57BL/6j wild-type mice were used for flow cytometry. Furthermore, lungs from Balb/c mice were collected for qRT-PCR, scRNAseq, and IHC & IF staining. Time points and pre-treatment is given in the respective figure/figure legend. For flow cytometry, single cells suspension of spleen was prepared by meshing the organ through a 70µm cell strainer (BD Falcon) and a subsequent lysis of erythrocytes in a hypotonic buffer. Single cell suspension of collected lungs was prepared for flow cytometry and scRNAseq. Lungs were digested enzymatically using Lung Dissociation Kit, mouse and gentleMACS^TM^ dissociator (Miltenyi Biotec) according to the manufacturer’s manual. Erythrocytes were lysed as described before. Lungs for qRT-PCR were collected 6 hours after the fourth RNA-LPX injection d13 (CT26-metastasis-bearing lungs were injected with RNA-LPX at d3, d6, d10 & d13). Lungs were transferred into liquid nitrogen directly after extraction and stored at −80°C upon use. Organs collected for IHC and IF staining were stored in 4% Histofix (Carl Roth) after extraction and embedded in paraffin.

### Flow Cytometry

Single cell suspensions of spleen or lung were stained with fixable viability dye (BD) for 15 min at 4°C. Cells were washed twice with PBS and Fc-block was added for 15 min. at 4°C. Extracellular Antibodies were mixed and diluted in Brilliant Stain buffer (BD). Cells were incubated with the antibody mastermix for 30 min at 4°C. Cells were washed once with PBS and permeabilized using FoxP3/Transcription Factor Staining Buffer Set (BD) according to the manufactureŕs instructions. Antibodies for intra-cellular targets were mixed and diluted in Perm Buffer (BD). Staining with the intracellular mastermix was performed for 30min at 4°C and the cells were washed twice with FACS buffer (PBS containing 5% FCS and 5 mM EDTA) afterwards. Samples were fixed in BD stabilizing fixative solution (BD) according to the manufacturer’s instructions.

For the staining of murine whole blood, blood was collected from either the retro orbital vein plexus or the vena fasciales (according to the approval of the local regulatory authority). The blood was transferred directly into a 96-well plate and undiluted extracellular antibody Mastermix and live dead dye (Thermo Fisher) were added. After 30min incubation at 4°C red blood cells were lysed adding diluted 10x lysing solution (BD) according to manufactureŕs instructions for 5 min at room temperature. Cells were washed twice with PBS and the intracellular staining was performed as described above.

Ex vivo stimulation and intracellular cytokine staining (ICS) was performed on splenocytes as well as on lung single cells in RPMI 1640 with 10% FCS, 1% MEM NEAA, 1% Sodium pyruvate 0.5% Pen/Strep and 0.1% 2-Mercaptoethanol. CD107a monoclonal antibody (BD), 0.05 µg/ml PMA and 1µg/ml Ionomycin were added. After 1h incubation 10µg/ml Brefeldin A and 0.2% Golgi Stop (BD) were pipetted into each well and cells were incubated for 5 h. Subsequent viability and extra cellular staining was performed as described above. For permeabilization BD Cytofix/Cytoperm™ Fixation/Permeabilization Solution Kit (BD) was used according to manufacturer’s instructions. Antibodies against intracellular targets as well as cytokines were mixed, diluted in Perm/Wash buffer (BD) and added to the permeabilized cells. After 30min incubation at 4°C cells were washed twice with Perm/Wash buffer (BD).

For flow cytometry of immune cells in BALF, the collected cell pellet was incubated for 5 min in a hypotonic buffer to lyse potential erythrocyte contamination. After washing in 1 ml FACS buffer, the cells were resuspended in an antibody master mix containing Fc-block and incubated for 40 min at 4 °C in the dark. Cells were washed twice with PBS and stained with a viability dye (Thermo Fischer) for 15 min at 4 °C in dark. Subsequent to washing in PBS, cells were fixed in BD stabilizing fixative solution (BD) according to the manufacturer’s instructions. Before acquisition TrueCount Beads (Thermo Fisher) were added for quantification.

Following monoclonal antibodies against extra- and intra cellular targets were used for FACS staining: 4-1BB, CD3, CD4, CD8, CD11b, CD11c, CD25, CD39, CD44, CD45, CD64, CD69, CD73, CD86, CD103, CD107a, CD127, Cleaved caspase3, Eomes, Epcam, F4-80, Foxp3, Gata3, Gr-1, Granzyme B, I-A/I-E, ICOS, IFN-γ, IL-2, Ki-67, KLRG-1, Ly6G, MHC II, NKp46, RORγT, TNF-a, XCR-1 and ordered from BD, Biolegend and Invitrogen.

For quantification of Immune cell subsets, samples were taken up in a solution with TrueCount Beads (Thermo Fisher) before acquisition. Calculation of cell counts was performed as described by the manufacturer.

Data acquisition was done at the LSRFortessa machine (BD), and data were analyzed using FlowJo software.

### Single cell RNA sequencing (scRNAseq)

Single cell suspension from tumor-bearing lungs were obtained as described in “tissue preparation”. 1 x 10^6^ cells per mouse per group (n=5 per group) were pooled and stained with a live-dead dye and CD45. These 5 x 10^6^ cells from either cytokine RNA mix or irrelevant RNA-treated metastasis-bearing murine lungs. Subsequently, cells were sorted in order to obtain viable CD45+ cells per group. Single cell transcriptome profiling was performed using the Chromium Next GEM Single Cell 3’ v3.1 (10X Genomics) kit, following the manufactureŕs instructions. Briefly, ∼10.000 sorted single-cell suspensions per sample were loaded in droplets and then barcoded cDNA libraries were generated. The libraries were then sequenced on an Illumina NovaSeqTM 6000 with the following read lengths: 28 cycles (Read 1), 8 cycles (i7 Index) and 91 cycles (Read 2).

*Processing of scRNAseq raw data*: The 10x Genomics Cell Ranger single-cell pipeline (v5.0.0) was used to process the raw sequencing data and to generate gene expression matrices for each sample. Reads were aligned to the mouse reference genome (refdata-gex-mm10-1.2.0). For downstream analysis and visualization, the Seurat (v4.1.1 and v4.2.0) [48] and SCpubr (v2.0.2) [49] R toolkits (with R version 4.1.0) were utilized.

*Cell QC, filtering and clustering scRNAseq:* Cells with fewer than 200 genes, or with gene or mitochondrial counts exceeding three median absolute deviation (MADs) above the median, were considered of low quality and excluded from further analysis. A total of 14,612 cells were analyzed: 7,772 from the cytokine RNA mix group and 6,840 from the irrelevant RNA control group. Following quality control, the counts were log normalized and 2,000 variable features selected for the integrative analysis. The samples were integrated using the “fastMNN” function included in the batchelor (v1.10.0) package [50]. Corrected counts were scaled and served as input for Principal component analysis (PCA). Next, shared nearest neighbor graph-based construction was performed using the top 30 principal components. Cell clusters were identified using the Louvain algorithm, implemented in the Seurat’s “FindCluster” function, and the chosen resolution of 1.6. A two-dimensional non-linear embedding of the cells was generated on the same PC components using UMAP for visualization of the clusters.

*Cell type annotation scRNAseq:* Cell types were annotated based on the differentially top expressed genes per cluster (identified by the Seurat’s “FindAllMarkers” function) and the comparison with canonical markers for immune cell types. Cell clusters with relatively lower RNA/gene content and no significantly (log2 fold change >0.5 and/or expressed in less than 25% of the cell population) expressed markers was assigned as “low quality clusters” and therefore removed for downstream analyses. Our approach defined 22 distinct cell clusters were : CD4-1, CD4-2, CD4-3, CD4+ Tregs-1, CD4+ Tregs-2, CD8-1, CD8-2, CD8-3, double-positive T cells (DPT cells), proliferating double-negative T (DNT) cells, B cells, NK1, NK2, NK3, NK-like cells, DC1/pDC, mast cells/DC2, pDC, MΦ, AMΦ, IMΦ, neutrophils. Cell subpopulation proportions, calculated as the percentage of the cells for each subpopulation over the total number of cells, have been compared between the two experimental groups and with FACS results.

### Differential gene expression analysis

Differential gene expression analysis between cytokine RNA mix and irrelevant RNA conditions were calculated, for each subset of cells forming a given immune cell type, using the Seurat’s “FindMarkers” function. Differentially expressed genes were then selected based on their adjusted p values (< 0.05).

### Cytokine Supernatant generation for treatment of human tumor primary cultures

Cytokines were produced in HEK293T/17 (ATCC). 1.2×106 cells/ well were seeded in a 6 well plate and let adhere overnight. The day after cells were transfected with 3 µg of total mRNA (HsIL2, hIFNa2a, HsIL7 in a 4:1:4 ratio) and 14.4 µl of Ribojuice (Merck, TR-1013). 20 hours after lipofection supernatants were collected and centrifuged (300xg, 4 min, RT), aliquoted in 5 mL aliquots and stored at −20 °C. As a control, supernatant of not transfected cells was used.

### Ex-vivo Human Tumor Primary Cultures

Punches of regular sizes (1.2 mm of diameter) were generated with a biopsy needle from the original tumor (WPI, WP1212). Punches were cultured in 80% organoid media composed of ADMEM/F12 (Gibco, 12634-010), 50% conditioned media Wnt/r-spondin/noggin (ATCC), 1mM HEPES (Invitrogen,15630080), Glutamax (1x, Invitrogen, 35050061), Nicotinamide 10mM (Sigma, 72340-250G), N-Acetilcysteine 1 mM (Sigma, A7250-25G), B-27 w/o vitamin A (1x, Invitrogen, 12587010), Pen/Strep Glutamine (1x, Invitrogen, 10378016), Gastrin (10 nM, Sigma, G9020-.5MG), SB-202190 (10 µM, Sigma, S7067-5MG), A83-01 (0.5 µM, Tocris, 2939), EGF (50 ng/ml, Invitrogen, PHG0311L) and Normocin (1x, InVivoGen, Ant-nr-1) and 20% of either control media or cytokine media. Each punch was distributed in one well of a 96-well plate (Ultra low attachment, Corning, 7007) and cultivated for 48 hours at 37 °C and 5% CO2. Punches were digested with 75 µg/ml Liberase DH (Roche, 5401054001) in ADMEM/F12 (Gibco, 12634-010) and 5 µg/ml DNAse (Sigma, 11284932001) for 60 min at 37°C. For immunohistochemistry, punches were fixed in Histofix (4%, Carl Roth, P087.3). Supernatant was collected and stored at −80°C for Codeplex analysis.

### Codeplex analysis (human primary tissue culture)

To measure GrzB from human cell culture supernatants, collected frozen supernatants were thawn and loaded to CodePlex Secretome Adaptive Immune-L Chips human (Isoplexis) according to the manufacturer’s protocol. Chips were installed together with CodePlex Secretome Adaptive Immune-L Panel human components (Isoplexis) into the IsoLight System (Isoplexis). To run the assay “Adaptive Immunsystem, CodePlex Secretome” program was used (Version 1.10 Isolight). Quantitative Data analysis was performed using IsoSpeak software version 2.9.

### Histology

Tumor tissues were harvested from mice, fixed with 4 % histofix overnight at 4 °C and embedded in paraffin blocks (FFPE). Human PDOs were fixed in 4% histofix 1 h at 4 °C and embedded in paraffin blocks (FFPE). 3 μm FFPE tissue sections were stained with haematoxylin and eosin (H&E). Briefly, after deparaffinisation and rehydration, sections were incubated in Hemelaum (acc. to Mayer) solution for five minutes and counterstained with eosin for two minutes. Light microscope images were taken using the Zeiss Axioscanner.

### Masson’s Trichrom staining (mouse)

3 µm FFPE tissue sections were deparaffinized and stained with different reagents in given order and incubation times, and washed in between with distilled water; 5 min in Hematoxylin (acc. to Harris (Merk 1.09253.0500)), 5 min in Mallory red (1 g acid fuchsin, 0,4 g Orange G in 300 mL distilled water and 1 mL acetic acid (100%)), 15 min in 1% phosphomolybdic acid solution, and counterstaining 5 min in light green solution (1 g light green + 1 mL acetic acid (100%) in 100 mL distilled water). Sections were dehydrated with Acetic acid – Ethanol (1:100), 100% Ethanol and Xylene. Images were acquired using Aperio AT2 Slide Scanner (Leica Biosystems) and analyzed with ImageScope software (Version 12.4.3.5008, Leica Bioscience).

### Immunohistochemistry (mouse)

Three-μm FFPE tissue sections were stained with rabbit polyclonal anti-Luc (Abcam, ab185924, 1:500) and anti-CD31 (Thermo Scientific, RB-10333, 1:200) antibodies. Briefly, after antigen retrieval using Sodium citrate buffer (10 mM, pH=6) for 10 min at 120°C in DAKO Pascal pressure chamber, primary antibody incubation was performed o/n at 4 °C. Next day, sections were washed in PBS, then incubated in HRP-anti-rabbit antibody (HRP anti-rabbit, Immunologic, DPVR-110HRP, RTU) 30 min at RT. The staining was detected using a streptavidin-HRP system (Vector Labs). Counterstaining was performed with Hemalaum (Carl Roth, T865.2) for 1min.

For manual 2 colour Opal mIHC staining, sections were first incubated with anti-Luc antibody (1:1000) 1 h at RT. After HRP-conjugated secondary antibody incubation, signal amplification was done using Opal 570 10 min at RT (1:100, Akoya Bioscience, FP1488001). A second antigen retrieval step was performed to remove first antibody-secondary antibody complex. Sections were then incubated with anti-F4/80 (Invitrogen, MA5-16363, 1:1000) antibody o/n at 4 °C. Opal 520 10 min at RT (1:100, Akoya Bioscience, FP1487001) was used for signal amplification after secondary antibody incubation. Counterstaining was performed with DAPI for 5min at RT (2 drop/1000ul), Akoya Bioscience, FP1490). Microscope images were taken using the Zeiss AxioImager M2 and analyzed using ZEN 2.3 (blue edition) software.

For a manual 4-colour Opal mIHC staining tissue sections were incubated with anti-anti-F4/80 (CST, 70076S, 1:1000) antibody o/n at 4°C. After HRP-conjugated secondary antibody incubation (Akoya Bioscience, DPVR110HRP), signal amplification was performed using Opal 480 (Akoya Bioscience, FP1500001KT, 1:150) for 10 min at RT. After another antigen retrieval step, samples were stained with anti-luciferase (Abcam, ab185924, 1:250) for 1h at RT followed by secondary anti-rabbit-HRP antibody staining for 30 min at RT and signal amplification using Opal 520 for 10 min at RT (Akoya Bioscience, FP1487001KT,1:250). A third antigen retrieval step was followed by anti-cytokeratin staining (DAKO, GA053, ready to use) o/n at 4°C, secondary antibody staining using MOM-HRP Detector (Abcam, MOM Polymer IHC Kit, ab269452) for 30 min at RT and signal amplification using Opal 570 (Akoya Bioscience, FP1488001KT,1:250) for 10 min at RT. After a last antigen retrieval step, samples were incubated with anti-CD31 (CST, 77699S, 1:500) for 1 h at RT. Secondary antibody staining was performed using anti-rabbit-HRP for 30 min at RT followed by signal amplification using Opal 690 (Akoya Bioscience, FP1497001KT,1:150) for 10 min at RT. Counterstaining was performed using DAPI for 5 min at RT (Akoya Bioscience, FP1490, 8 drops/1ml). Images were acquired using the PhenoImager HT multispectral slide scanner (Akoya Biosciences) and analyzed using Phenochart (Version 1.1.0) software.

For automated Opal mIHC staining of human PDOs, 3 μm FFPE tissue sections were stained in the Leica BOND Rx autostainer using a rabbit anti-CD8 (Clone: SP16) antibody (1:100, Novus Biologicals, NBP2-26484), a rabbit anti-Ki67 (Clone: SP6) antibody (1:500, Thermo Scientific, RM-9106), a mouse anti-Granzyme B (Clone: 11F1) antibody (RTU, Leica Biosystems, PA0291) in the given order. The standard mIHC protocol adjusted to 4-colour with 30 min primary antibody incubation each was used. Microscope images were taken using the PhenoImager HT multispectral slide scanner (Akoya Biosciences) and analyzed using Phenochart (Version 1.1.0) software.

### RNA In Situ Hybridization

ISH was performed using custom made Luc probe (Advanced Cell Diagnostics) and RNAscope detection reagents 2.5 HD Brown (Advanced Cell Diagnostics, 322000) according to the manufacturer-supplied protocol. Standard pre-treatment was done using 1x Target retrieval buffer for 15min at 95°C (Advanced Cell Diagnostics, 322330) and Protease plus for 30min at 40°C. (Advanced Cell Diagnostics, 322331). Counterstaining was performed with Hemalum (Carl Roth, T865.2) for 1min. Light microscope images were taken using the Zeiss Axioscanner.

### Phenocycler Fusion assay

The PhenoCycler®-Fusion system (Akoya Biosciences) was used to analyze T cells within tumor microenvironment. Sample preparation and tissue staining were performed according to the PhenoCycler Fusion user manual (Akoya, PhenoImager Fusion SW Version: 2.1.0). Briefly, 3 μm FFPE tissue sections were pre-processed by deparaffinization, dewaxing/rehydration, and heat-mediated antigen retrieval (Akoya, 7000017). Then, tissues were simultaneously stained with the entire barcoded-antibody panel (see table below). After the staining, a flow cell (Akoya, 240205) was placed on top of the slide.

For the reporter plate, unique and spectrally distinct reporters, complementary to the barcodes used in the antibody panel, were organized into groups of three and combined with a nuclear stain (Akoya, 7000003). Each group constituted a separate PhenoCycler cycle. The experimental protocol and reporter plate design were carried out using the PhenoCycler Experiment Designer software. The PhenoCycler run was fully automated and executed by the Controller software. PhenoCycler reporters were delivered to the tissue by the PhenoCycler instrument and detected using the Fusion slide scanner. The repetition of these cycles with different reporters allowed the visualization of the complete antibody panel on the same tissue area.

| Antibody | Cat.No. | Dilution | Fluorophore | Cat.No. | Dilution | Cycle |
| --- | --- | --- | --- | --- | --- | --- |
| CD8-BX026 | 200092 | 1:200 | RX026-ATTO550 | 6250007 | 1:50 | 1 |
| Lag3-BX001 | Custom | 1:50 | RX001-AF750 | 6450001 | 1:50 | 1 |
| CD4-BX003 | 200077 | 1:200 | RX003-AF647 | 6550025 | 1:50 | 1 |
| CD3e-BX080 | 200075 | 1:200 | RX080-AF647 | 6550031 | 1:50 | 2 |
| TOX-BX060 | 240126 | 1:200 | RX060-ATTO550 | 6250023 | 1:50 | 2 |

### Image analysis Phenocycler Fusion assay

Quantification of luciferase expressing cell types and CD8 T cell exhaustion were performed using Visiopharm software (Version 2025.02.2.18022). Tissue was detected based on DAPI signal and cells were segmented using the Cell Detection AI (Fluorescence) Analysis Protocol Package (Version 2023.01.1.13576) developed by Visopharm. Cells were phenotyped using the thresholding function of the Phenoplex workflow.To identify luciferase expressing cell types, cells were thresholded based on cellular positivity for CD31, cytokeratin, F4/80 and luciferase, respectively. To quantify CD8 T cell exhaustion, cells were thresholded based on their cellular expression of CD8 and Lag3 as well as the nuclear expression of Tox. Tissue regions with an unspecific background in the Tox channel were excluded from analysis.

Visiopharm data was further processed using R studio (Version 2024.04.0) and Microsoft Excel (Microsoft 365 MSO, Version 2507) to analyse the cell type distribution of luciferase-positive cells or the percentage of exhausted CD8 T cells. Miss-classified cells with unclear phenotypes were excluded from analysis.

### Design and validation of Primers for qRT-PCR

Primers used in quantitative Real-Time PCR (qRT-PCR) were designed to bind specifically mRNA-related transcripts of the target genes of interest, assisted by “Oligo Analysis Tool” (https://eurofinsgenomics.eu/en/ecom/tools/oligo-analysis/), “OligoAnalyzer” (https://eu.idtdna.com/calc/analyzer), “Primer-BLAST” (https://www.ncbi.nlm.nih.gov/tools/primer-blast/), “UCSC In-Silico-PCR” (https://genome.ucsc.edu/cgi-bin/hgPcr) and “UNAFold - mFold DNA Folding Form” (http://www.unafold.org/mfold/applications/dna-folding-form.php).

The assays were validated using a BioRad CFX384 Touch Real-Time PCR Detection System and BioRad’s SsoAdvanced Universal SYBR Green Supermix determining the optimal annealing temperature by comparative runs, including melting curve analysis. Correct PCR product size was controlled by capillary gel electrophoresis (QIAGEN QIAxcel or Agilent Fragment Analyzer). Primer efficiencies, limit of quantification (LOQ) and limit of detection (LOD) were determined using dilution series.

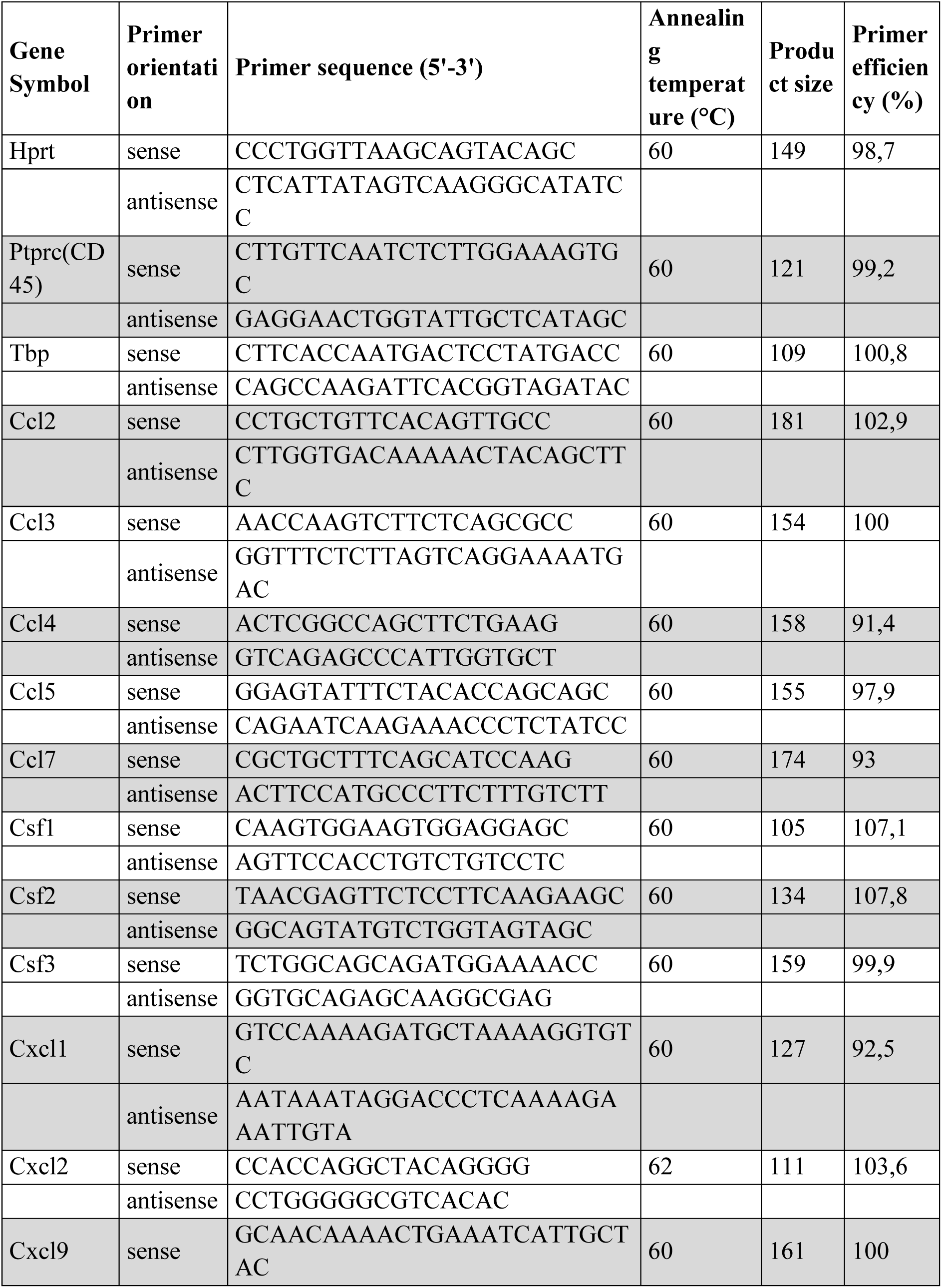
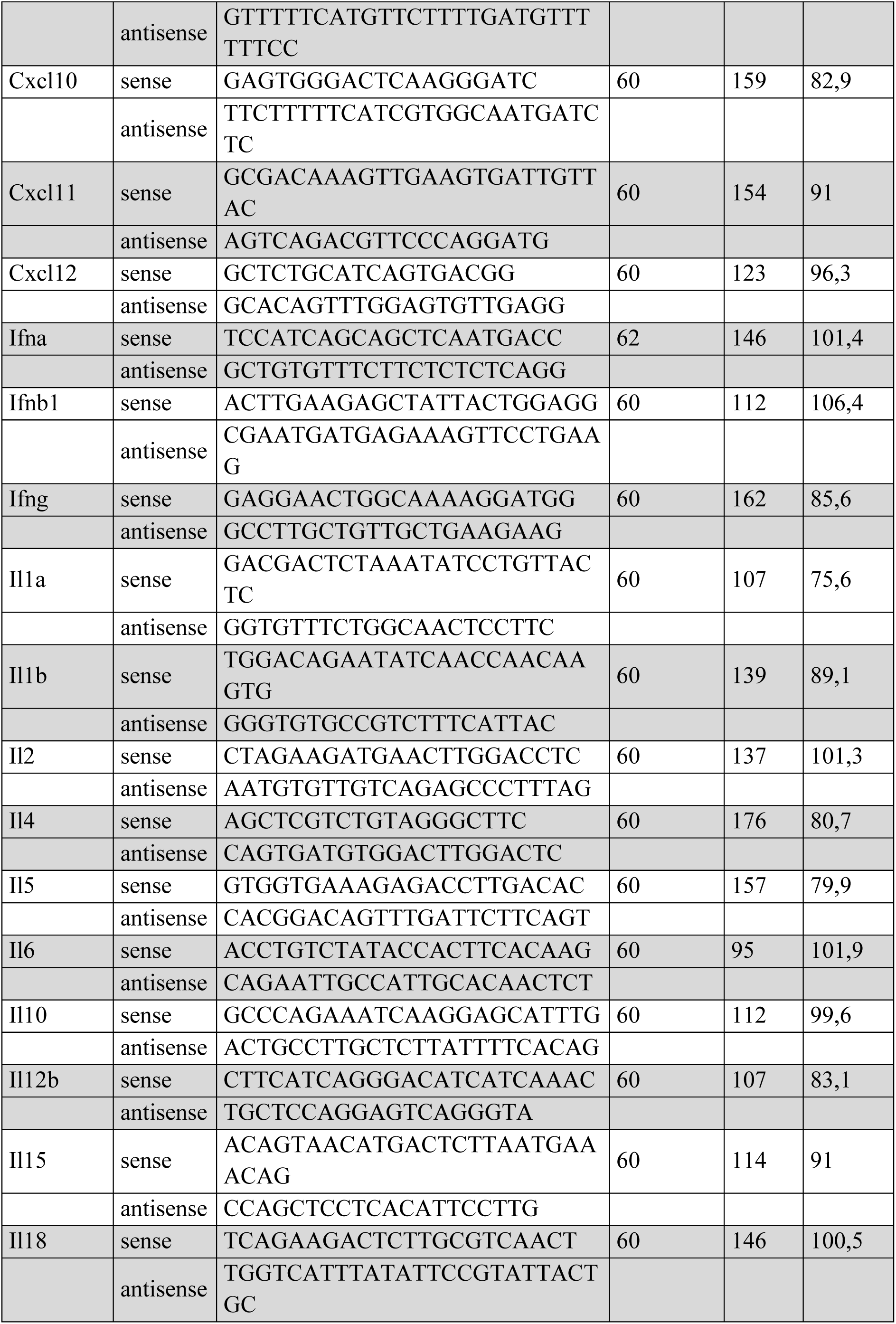
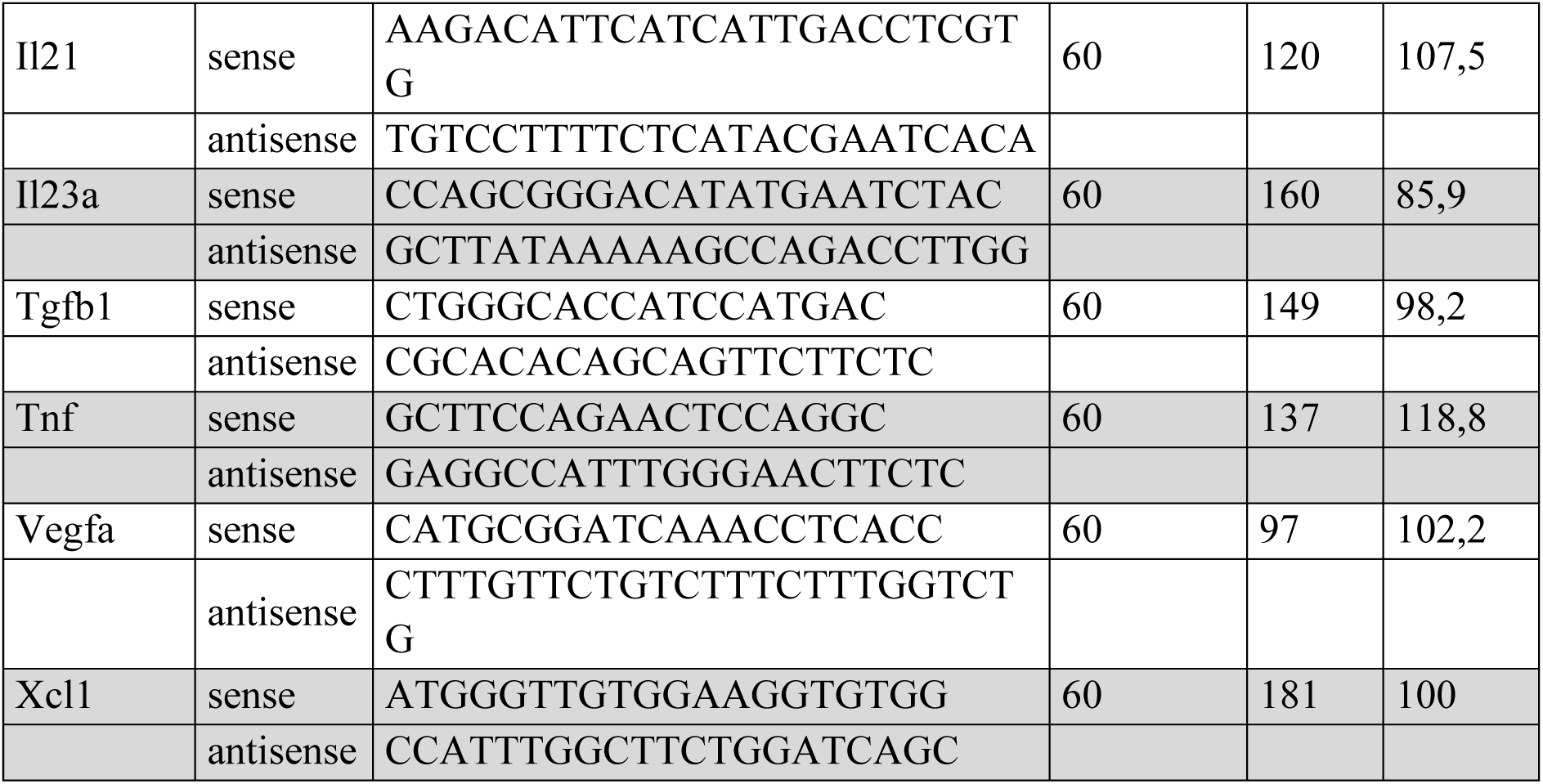

### Sample preparation and nucleic acid extraction for qRT-PCR

Mouse lung tissues were received as fresh frozen (FF) samples. Using phenol/chloroform extraction, Precellys Evolution Touch Homogenizer (bertin technologies) and QIAGEN’s RNeasy Mini Kit, following the manufacturer’s protocol, total RNA was extracted from the samples. An on-column DNase I digestion was performed. RNA concentration and purity was determined via ThermoFisher NanoDrop 2000c UV/VIS spectrometry instrument, RNA integrity was controlled via capillary gel electrophoresis (Agilent Fragment Analyzer).

### Reverse Transcription

Extracted mouse total RNA was reverse transcribed to cDNA using TAKARA PrimeScript RT Reagent Kit using 3 µg total RNA per reaction, following the manufacturer’s protocol, including the gDNA Eraser step. Each 60 µL RT reaction was finally diluted 1:3 with 120 µL of nuclease-free water.

### qRT-PCR on mouse lung samples

Quantitative Real-Time PCR was performed on a BioRad CFX384 Touch Real-Time PCR Detection System. All reactions were run in three technical replicates. As positive controls mouse reference RNA (Agilent), naïve and concanavalin A stimulated mouse splenocytes were used. Water was used as NTC and total RNA without reverse transcription served as NAC. Each PCR had an overall volume of 15 µL, containing 7.5 µL SsoAdvanced Universal SYBR Green Supermix (BioRad), 5.25 µL PCR-grade water and 0.5 µL of 10 µM sense and antisense primer, resulting in a final concentration of 333 nM per primer. The thermal cycle protocol was run with an initial heat activation and denaturation at 98°C for 30 sec, and 40 cycles of two-step PCR with a denaturation step at 98°C for 10 sec and a combined annealing & elongation step for 30 sec at 60°C or 62°C (see primer list). Finally, a melting curve analysis was performed using the instrument’s standard parameters. Cq-values were determined using regression mode in BioRad CFX Manager (Version 3.1.1517.0823).

### Analysis and Results of qRT-PCR data

For expression analysis the median Cq-value of each technical triplicate was used and normalized to the median reference gene expression (Hprt and Tbp) using ΔΔCq-calculation. Primer efficiencies were considered in the determination of the expression values. Depicted fold changes resulted from comparing the median expression +1 of each individual biological replicate in the treatment group (cytokine RNA mix) versus the median relative expression +1 of all biological replicates of the control group (irrelevant RNA).

### Statistical analysis

Statistical analysis was performed using GraphPad Prism 7 or 8 software (Graphpad Software, Inc.), except the weighed logrank test (see Fig. 5B,C) where testing was performed with “R” under full implementation of the Fleming-Harrington class for right-censored data. Specific tests are indicated in the figure legends; typically, significance in the survival experiments was determined via Mantel-Cox logrank test, pairwise comparisons were performed unpaired two-tailed t-test, and ordinary one-way ANOVA with post-hoc Tukey’s test were used for multiple comparisons. Values of p ≤ 0.05 were considered to be statistically significant; *p ≤ 0.05, **p ≤ 0.01, ***p ≤ 0.001, ****p ≤ 0.0001. Error bars indicate standard error of the mean (SEM).

### Data and materials availability

Further information, (scRNAseq) raw data and material may be requested from the authors and will be available upon reasonable interest.

## LIST OF SUPPLEMENTARY MATERIAL

Supplementary Fig. S1 to S10

## Acknowledgments

Evelyn Russel, Magdalena Brkic, Ines Beulshausen, Anindhita Muralidharan, Alekhya Porapu, Steven Zenner, Ute Schmitt, Evelin Petscherskich, Kimberley Wohde, Michelle Jäger, Münteha Yilmaz, Jessica Sriha, Anh-Phi Duong for excellent technical assistance; Martin Kapp, Miroslav Dörr, Larry Kwesi Sarpong, Ehsan Mehravar for providing formulations, Abderraouf Selmi and Maik Schork for supporting experiments, Ioanna Tsoukala for flow cytometry of BALF, Julia Zimmermann for Phenocycler Fusion staining and quantification, Ann-Kathrin Koch, Anne Kölsch and Yasemin Ahrberg for supporting the tumor punch platform, Alessandra Gargano for supporting IF experiments, Krutika Khinvasara for supporting scRNAseq, Nina Köhl for supporting qRT-PCR Assays, Sebastian Attig for cell sorting, Sonja Witzel and Bodo Tillman for cloning, Tommaso Torcellan, Olga Ucar & Fiona Powel for copyediting the manuscript. Thomas Rösler for weighted logrank test. Kurt Reifenberg for excellent support as animal welfare officer, Jan Beck and Mathias Vormehr for generating CT26 gp70 k.o. and B2M k.o. cell lines.

## Author contributions

A.K., F.V., M.D. and U.S. conceptualized the work. A.K., F.V., S.L., planned and analyzed experiments. S.L. and A.K. performed experiments, A.K., F.V., M.D., S.K., and U.S. interpreted the data. A.K. wrote the original draft, S.L., F.V., M.D edited the manuscript, M.S. designed, performed and analyzed qRT-PCR assays. E.D. and A.F. performed scRNAseq., E.F., A.F. and A.K. interpreted scRNAseq data, O.A.K. and M.M.G. provided clinical samples, performed histological and immunofluorescent stainings. S.F.K. and A.M. performed human *in vitro* Treg suppression assay, U.S., F.G. and A.M. established cytokine variants, E.S. planned & performed *ex vivo* human primary tissue culture. H.H., I.H.E.B. and F.B. optimized formulations, F.B. and K.T. provided formulations, S.F.K. and A.M. did pharmacokinetics analysis. J.B. and M.M. performed and analyzed PET experiments, J.P. and K.W. planned and analyzed experiments related to formulation characterization. All authors reviewed the manuscript and approved the final version of the manuscript for submission.

