## Supplementary Figures for "Lung-targeted cytokine-coding RNA-lipoplexes induce T and NK cell-mediated anti-tumor immune response"

**Overview:**

**Supplementary Figure S1:** Lung-targeting F5 formulation of surrogate RNA leads to protein production within the lung tissue as well as tumor tissue in the lung.

**Supplementary Figure S2:** Histological examination and BALF analysis show no signs of particle, nor cytokine-induced inflammation, vascular thrombosis, edema, or diffuse alveolar damage.

**Supplementary Figure S3:** Treatment with cytokine RNA mix induces activation of T cells and NK cells in the lungs and spleens of B16F10 metastasis-bearing mice.

**Supplementary Figure S4:** Cytokine RNA mix induced myeloid activation, increased ILC2/ILC3 numbers and reduced activation of $CD4^{+}Foxp3^{+}CD25^{+/-}$ cells in CT26 metastasis-bearing lungs as well as immunomodulatory effects in spleen and peripheral blood.

**Supplementary Figure S5:** Cluster definition and cell characteristics from scRNAseq analysis.

**Supplementary Figure S6:** Characteristics of $CD8^{+}$ T and NK cell clusters in scRNAseq analysis.

**Supplementary Figure S7:** NK cells are major producers of *Gzmb*, IFN-γ release is not playing a dominant role for antitumoral efficacy.

**Supplementary Figure S8:** Top differentially expressed genes from scRNAseq analyses.

**Supplementary Figure S9:** Efficacy of the cytokine RNA mix is depending on synergism of IFN-α with Alb-IL-2mut and IL-7-Alb.

**Supplementary Figure S10:** Cytokine RNA mix treatment does not majorly increase exhaustion and expression of co-inhibitory molecules and synergizes with chemotherapy.

Supplementary Figure S1

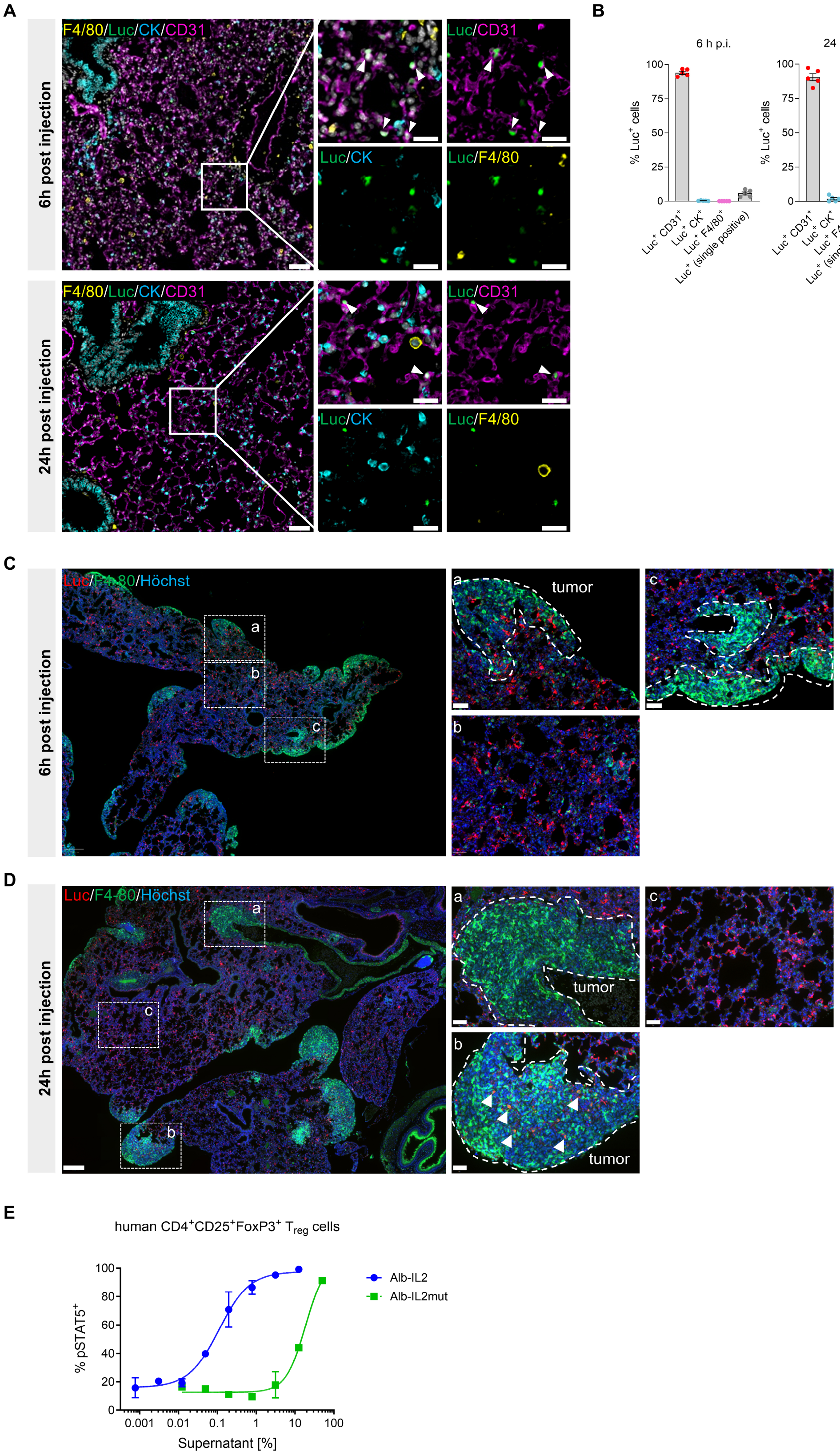

**Supplementary Figure S1. Lung-targeting F5 formulation of surrogate RNA leads to protein production within the lung tissue as well as tumor tissue in the lung.** (**A, B**) BALB/c mice (n=5) were i.v. injected with lung-targeting RNA-LPX containing 10 µg luciferase (luc)-encoding RNA and sacrificed 6 or 24 hours after RNA-LPX injection. (A) Representative immunofluorescence staining of mouse lung showing Luc (green), F4-80 (yellow, macrophages), cytokeratin (CK, turquoise, epithelial cells), and CD31 (red, endothelial cells), DAPI/nucleus (grey, nucleus) 6 hours after injection (**B**) Quantification of luciferase-transgene expressing cells at 6 or 24 hours post injection (n=5). (**C**, **D**) 10 µg luciferase-coding RNA formulated as RNA-LPX was administered i.v. to CT26 metastasis-bearing BALB/c mice 10 days after tumor inoculation. Lung tissue was collected 6 (**C**) or 24 hours (**D**) after RNA-LPX administration (n=3 per time point). Immunofluorescent staining for Luc (red), F4/80 (green) and Hoechst (blue, nuclei) was performed (white dashed line indicates tumor tissue). Luc expression is observed throughout the lung tissue 6 and 24 h post injection and can be detected in some of the tumor tissue. Only some of the F4/80$^{+}$ macrophages (green) show Luc expression (white arrows). Scale bars: 250 µm and 50 µm. (**E**) Decreased $T_{reg}$ activation by Alb-IL-2mut compared with Alb-IL-2 (“wildtype”) as shown by STAT5 phosphorylation in an *in vitro* assay employing PBMCs treated with supernatants of RNA-encoded cytokine-lipofected HEK cells.

### Supplementary Figure S2

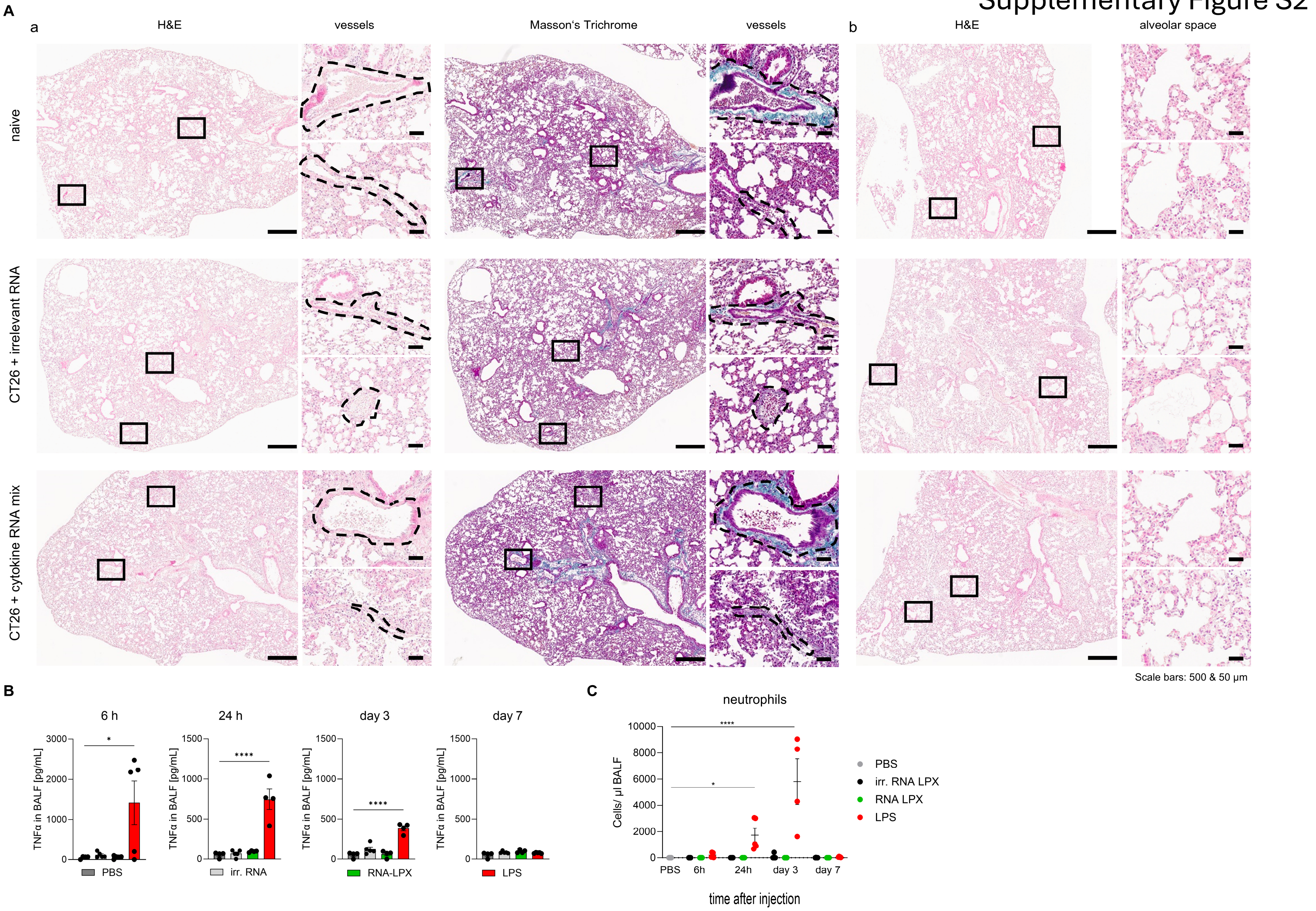

**Supplementary Figure S2. Histological examination and BALF analysis show no signs of particle, nor cytokine-induced inflammation, vascular thrombosis, edema, or diffuse alveolar damage.** (**A**) BALB/c mice (n=5) were injected with CT26 cells i.v., irrelevant RNA or cytokine RNA mix as LPX was administered twice per week for a total of 4 injections, or naïve BALB/c mice (n=5) were used as control. Mice were sacrificed 6 hours after the last injection and lungs were subjected to H&E or Masson's Trichrome staining Vessels (black dashed lines) are shown as zoom-ins (a) revealing no signs of perivascular infiltration typical for IL-2 mediated toxicity (H&E), no endothelial damage or general widespread lung inflammation. Masson's Trichrome staining shows no signs of particle-induced clotting or thrombosis. Zoom-ins to the alveolar space (b) show no signs of hyaline membranes, as a typical early arising feature of acute phase of acute respiratory distress syndrome (ARD) or diffuse alveolar damage (DAD) and no signs of pulmonary edema. Shown is 1 representative lung out of 5 mice with the vessels (black dashed lines) are shown as zoomed-in images. Scale bars: 500 µm and 50 µm. **(B**, **C**) BALB/c mice (n= 20 per group) were i.v. inoculated with either irrelevant RNA or cytokine RNA mix containing lung-targeting RNA-LPX (30µg total RNA/mouse) or with PBS (n=5 mice; negative control). An additional group of mice (n=20) was treated intranasally with 3 mg/kg LPS (positive control). Mice (n=5 per group) were sacrificed after 6, 24 hours, 3- or 7-days post treatment and BALF was collected. For animal welfare reasons (3R principles), the PBS –treated group was analyzed only once (at 6 hours), as no changes in BALF composition were expected, these values are therefore displayed across all timepoints. Of note, a total of 6 samples were excluded due to technical issues during sampling. (**B**) TNF-α in BALF as determined by ELISA. Statistical significance was determined via one-way ANOVA and Dunnett's multiple comparisons with PBS as control group. (**C**) Flow cytometric analysis of neutrophils in BALF displayed as total cells per µl of BALF. Statistical significance was determined via two-way ANOVA and Dunnett's multiple comparisons test with PBS as control group.

Supplementary Figure 3

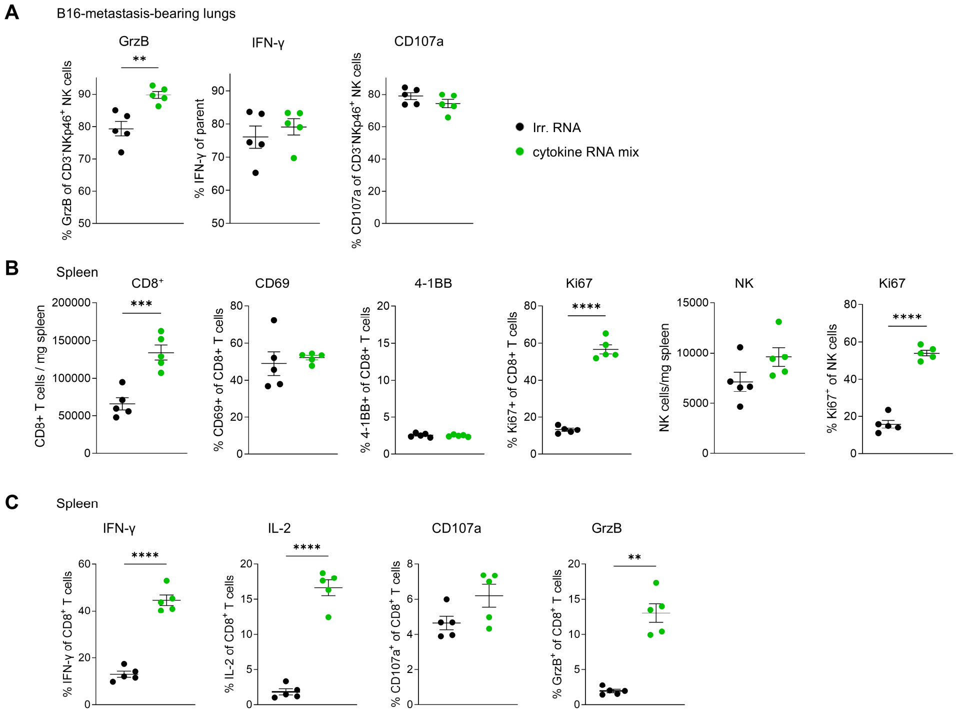


**Supplementary Figure S3. Treatment with cytokine RNA mix induces activation of T cells and NK cells in the lungs and spleens of B16F10 metastasis-bearing mice.** C57BL6 mice (n=5 per group) were injected with B16F10-*luc* tumor cells i.v. and treated with cytokine RNA mix or irrelevant RNA as shown in Fig. 2B. (**A**) GrzB, IFN-γ release and degranulation (CD107a) of NK cells from B16F10 metastasis-bearing lungs and stimulated *ex vivo* as shown in Fig. 2D. (**B**) Cell numbers per mg spleen weight and expression of activation markers in $CD8^+$ T cells and NK cells from spleens of B16F10 lung metastasis-bearing mice. (**C**) Cytokine and activation marker expression in $CD8^+$ splenocytes isolated from B16F10 lung metastasis-bearing mice after *ex vivo* stimulation. Statistical significance was determined using a two-tailed t-test for unpaired samples.

### Supplementary Figure S4

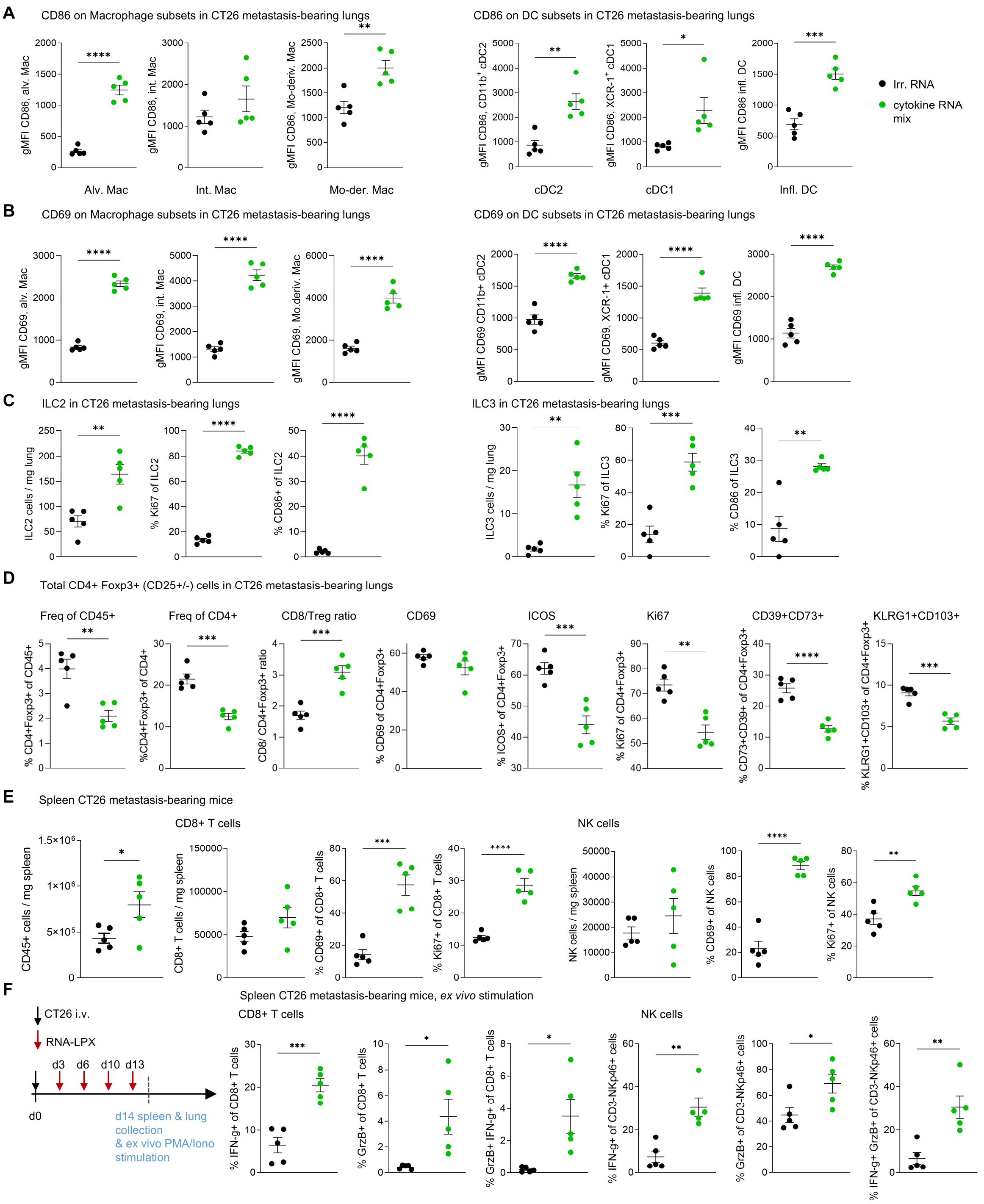

**Supplementary Figure S4. Cytokine RNA mix induced myeloid activation, increased ILC2/ILC3 numbers and reduced activation of $CD4^+Foxp3^+CD25^{+/-}$ cells in CT26 metastasis-bearing lungs as well as immunomodulatory effects in spleen.** (A-E) BALB/c mice (n=5 per group) were injected with CT26 tumor cells i.v. and treated with cytokine RNA mix or irrelevant RNA as described in Fig. 3D-F. Flow cytometric analysis of macrophage and DC subsets in metastasis-bearing lungs show increased activation (**A**) and CD86 expression (**B**) as quantified by geometric mean fluorescence intensity (gMFI). Alveolar Macrophages were gated as $CD64^+F4\text{-}80^+CD11c^{hi}MHCII^{low\text{-}int}$, interstitial macrophages as $CD64^+F4\text{-}80^+CD11c^-MHCII^{hi}$ and monocyte-derived macrophages as $CD64^+F4\text{-}80^+CD11c^{int}MHCII^{hi}$ .cDC2 were gated as $CD64^-F4\text{-}80^-CD11c^+MHCII^+CD11b^+$, cross-presenting cDC1 as $CD64^-F4\text{-}80^-CD11c^+MHCII^+CD11b^-$ $CD103^+XCR1^+$ , inflammatory DCs as $CD64^{int}F4\text{-}80^-CD11c^+MHCII^+$ $Gr\text{-}1^+CD11b^+$. (**C**) ILC2 and ILC3 in metastasis-bearing lungs were gated on non-T and non-classical NK ($NKp46^+Eomes^+$) on $CD127^{+/int}CD25^+$ cells and divided into ILC2 via Gata-3 and ILC3 via RORγt expression. Absolute numbers of ILC2 and ILC3 are increased as well as expression of Ki67 and CD86. (**D**) Since in lungs also a major population of $CD4^+Foxp3^+CD25^-$ $T_{reg}$ consists, $CD4^+Foxp3^+CD25^{+/-}$ cells in metastasis-bearing lungs were gated and absolute numbers, activation, proliferation, CD8/$T_{reg}$ ratio and phenotype are shown. (**E**) Flow cytometric analysis of CD8+ T cells and NK cells in spleen. (**F**) Functional analysis splenocytes: after tumor cell inoculation, mice were treated with RNA-LPX at d3, d6, d10, and d13 p.t.i., mice were sacrificed at d14 for an *ex vivo* stimulation assay with PMA/Ionomycin (as described in Fig. 3G). Significance for pairwise comparisons was determined by unpaired two-tailed t-test.

### Supplementary Figure S5

A

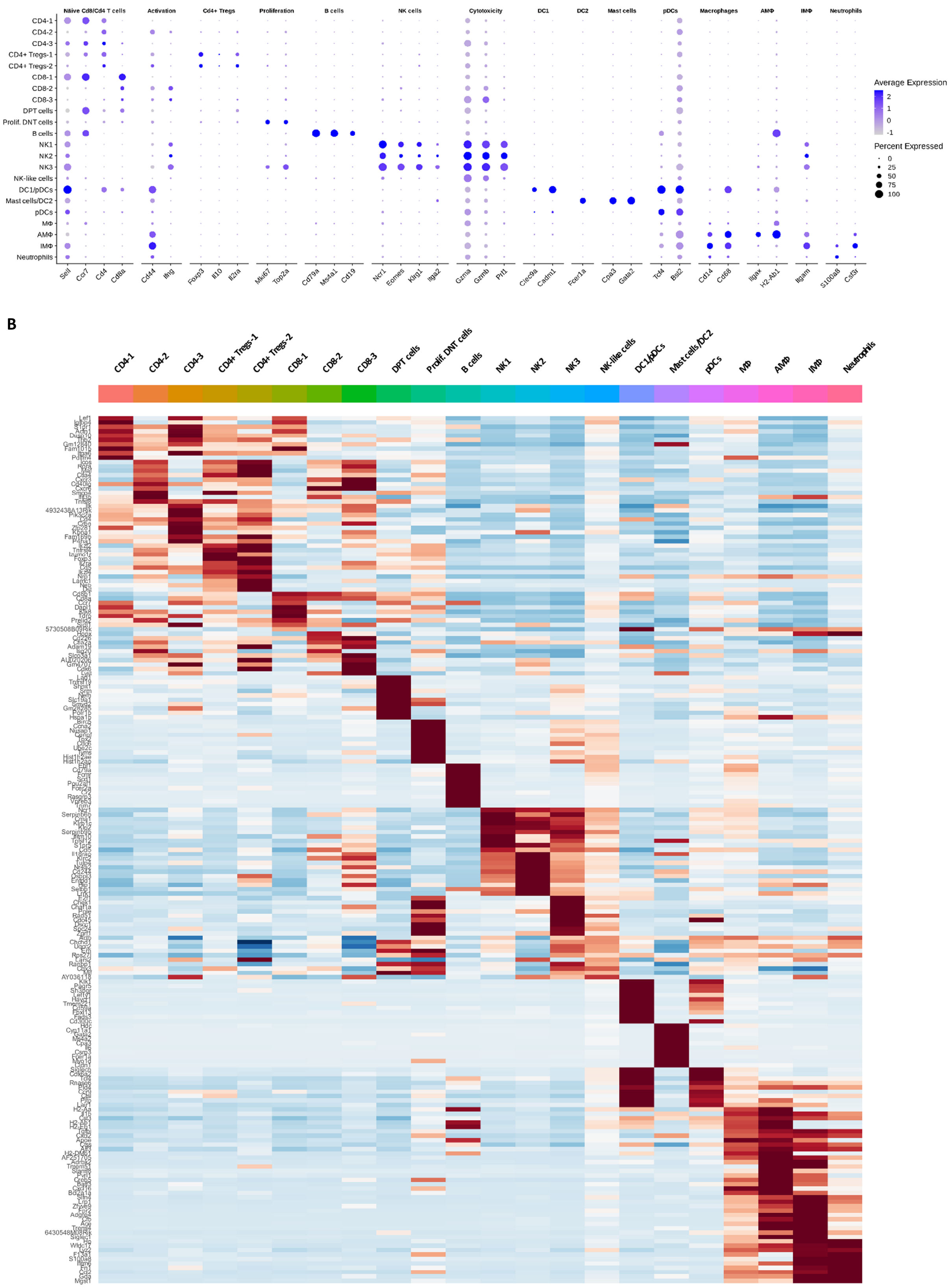

**Supplementary Figure S5. Cluster definition and cell characteristics from scRNAseq analysis.** BALB/c mice (n=5 per group) were i.v. injected with CT26 tumor cells, cytokine RNA mix or irrelevant RNA was i.v. administered as LPX at d3, d6 and d10 p.t.i., mice were sacrificed at d11, lungs were collected and pooled at same ratios before sorting of $CD45^+$ cells. $CD45^+$ cells were subjected to scRNAseq (related to Fig. 4). (**A**) Selected characteristics of 22 assigned clusters (**B**) Heatmap of the top 10 differentially expressed genes of each cell type population.

Supplementary Figure S6

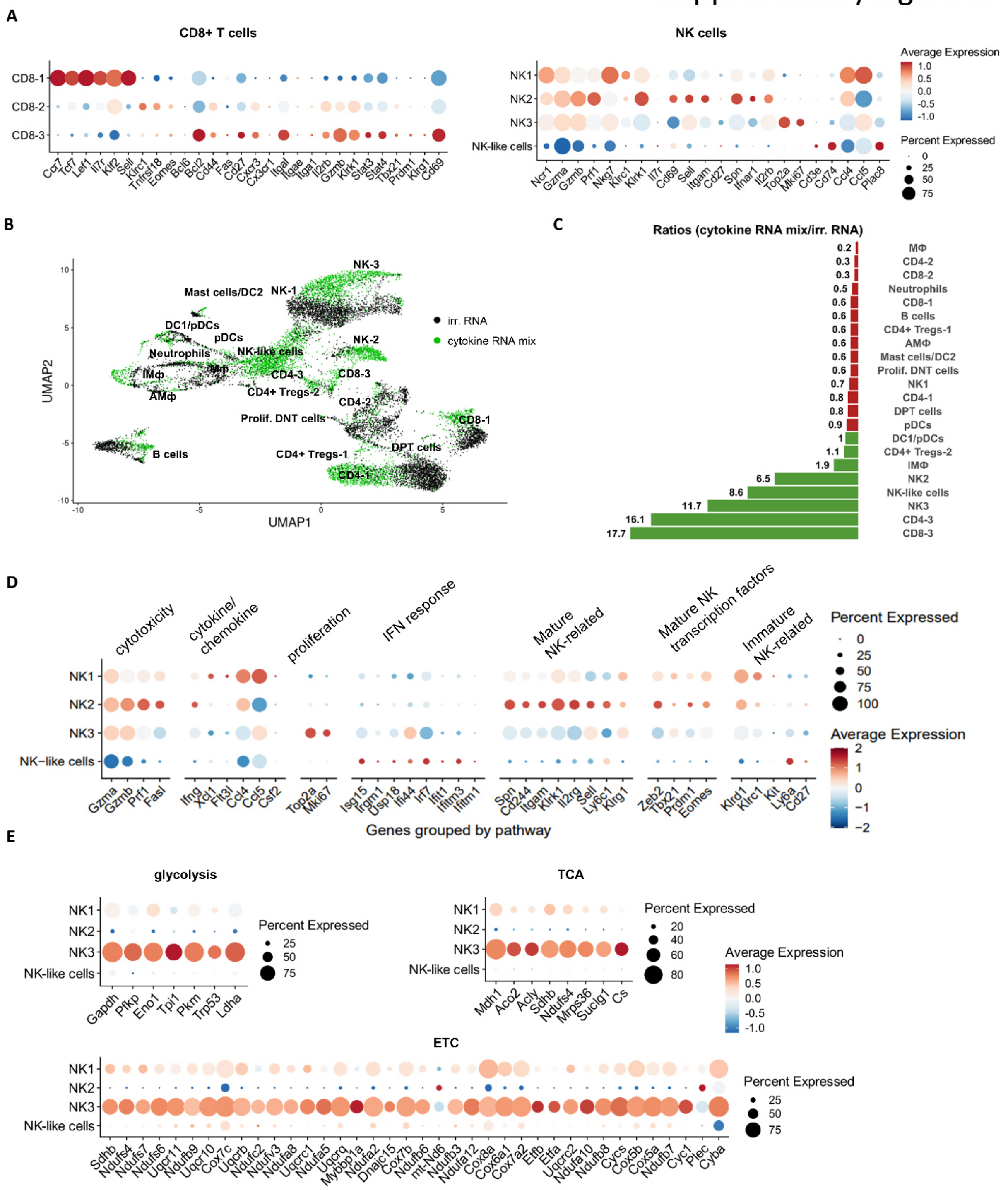

**Supplementary Figure S6. Characteristics of CD8$^+$ T and NK cell clusters in scRNAseq analysis.** BALB/c mice (n=5 per group) were i.v. injected with CT26 tumor cells, cytokine RNA mix or irrelevant RNA was i.v. administered as LPX at d3, d6 and d10 p.t.i., mice were sacrificed at d11, lungs were collected and pooled at same ratios before sorting of CD45$^+$ cells. CD45$^+$ cells were subjected to scRNAseq (related to Fig. 4). (**A**) Selected markers expressed on CD8+ T cells and NK cell subsets, (**B**) integrated UMAP of irrelevant RNA and cytokine RNA mix treated mice, (**C**) ratio of cytokine RNA mix-treated relative to irrelevant RNA-treated mice. (**D**) Selected maturation-related markers expressed on NK cell subsets (**E**) Selected metabolic genes expressed on NK cell subsets relevant for glycolysis, tricarboxylic acid cycle (TCA) or electron transport chain (ETC).

### Supplementary Figure S7

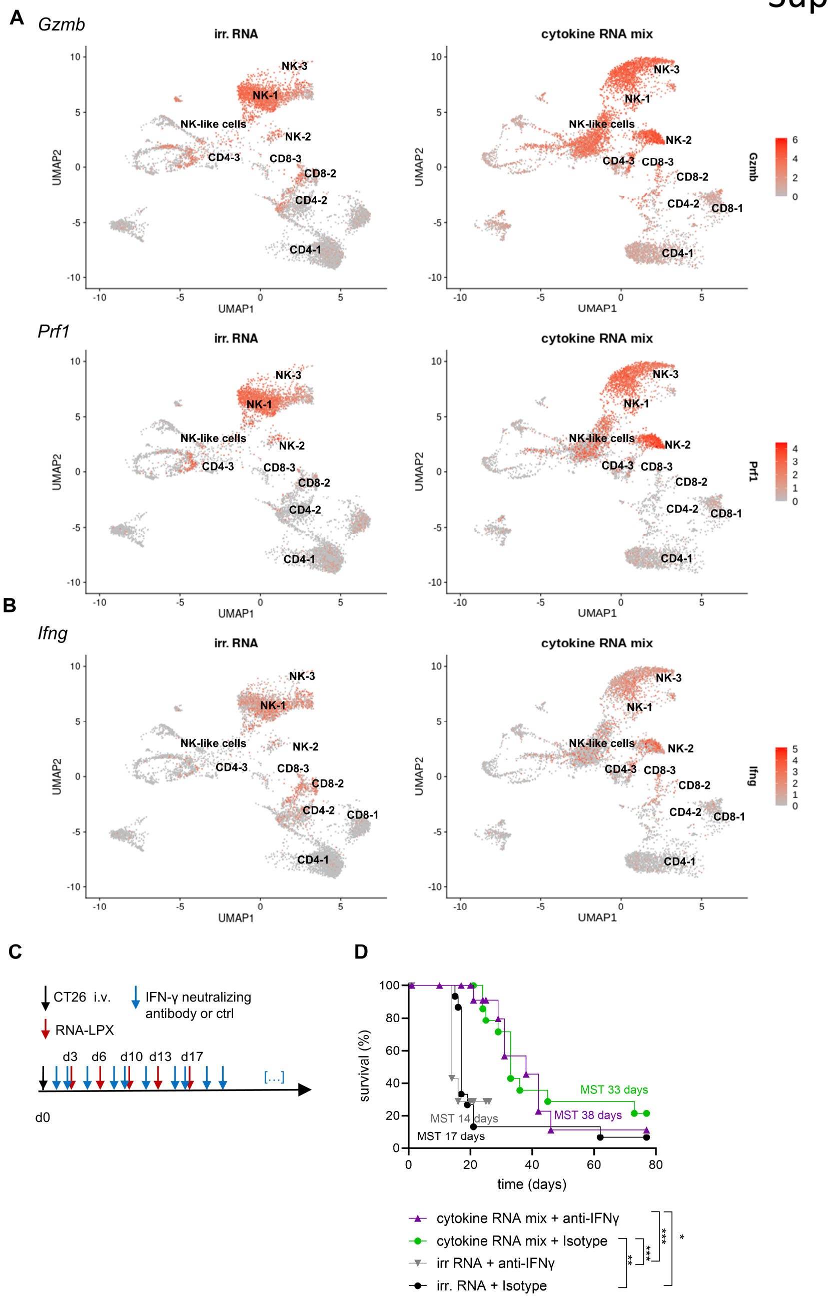


**Supplementary Figure S7. NK cells are major producers of *Gzmb*, IFN-γ release is not playing a dominant role for antitumoral efficacy.** (**A, B**) BALB/c mice (n=5 per group) were i.v. injected with CT26 tumor cells, cytokine RNA mix or irrelevant RNA was i.v. administered as LPX at d3, d6 and d10 p.t.i., mice were sacrificed at d11, lungs were collected and pooled at same ratios before sorting of $CD45^+$ cells. $CD45^+$ cells were subjected to scRNAseq (related to Fig. 4). Feature plots *Gzmb*, *Prf1* and *IFN-γ* comparing irrelevant RNA and cytokine RNA mix –treated mice from scRNAseq. (**C**) Experimental layout for (**D**); briefly: BALB/c mice (n = 15 per group) were injected with CT26 tumor cells i.v., cytokine RNA mix or irrelevant RNA were administered twice per week for a total of 5 injections as indicated and monitored until termination criteria were reached. IFN-γ-neutralizing antibody or isotype treatment was started 2 days prior to RNA-LPX treatment with an increased loading dose to ensure neutralisation before treatment start and was followed three times per week. Survival according to termination criteria is shown in (**D**). Survival was analyzed via Mantel-Cox logrank test.

### Supplementary Figure S8

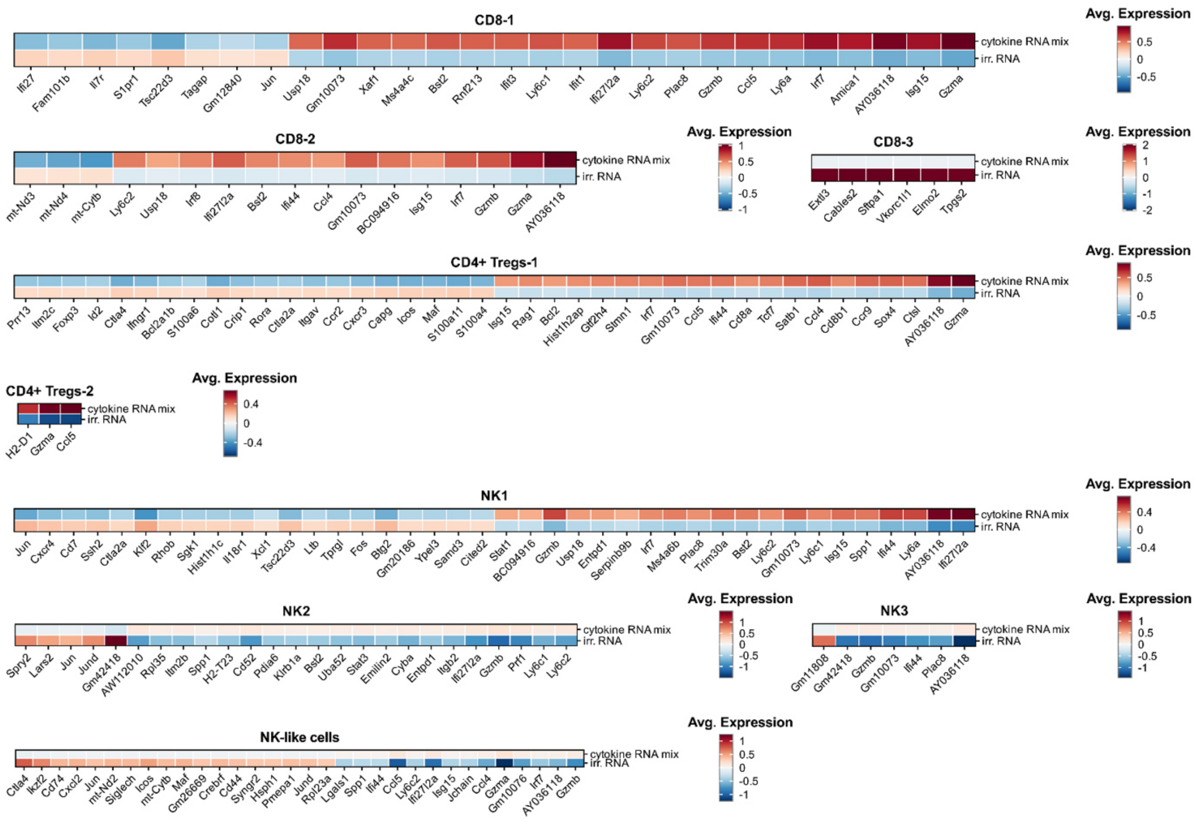

**Supplementary Figure S8: Top differentially expressed genes from scRNAseq analyses.** BALB/c mice (n=5 per group) were i.v. injected with CT26 tumor cells, cytokine RNA mix or irrelevant RNA was i.v. administered as LPX at d3, d6 and d10 p.t.i., mice were sacrificed at d11, lungs were collected and pooled at same ratios before sorting of $CD45^{+}$ cells. $CD45^{+}$ cells were subjected to scRNAseq (related to Fig. 4). Differential gene expression analysis between cytokine RNA mix and irrelevant RNA in CD8-1, CD8-2, CD8-3, $CD4^{+}$ Tregs-1, $CD4^{+}$ Tregs-2, NK1, NK2 and NK3, NK-like clusters.

#### Supplementary Figure 9

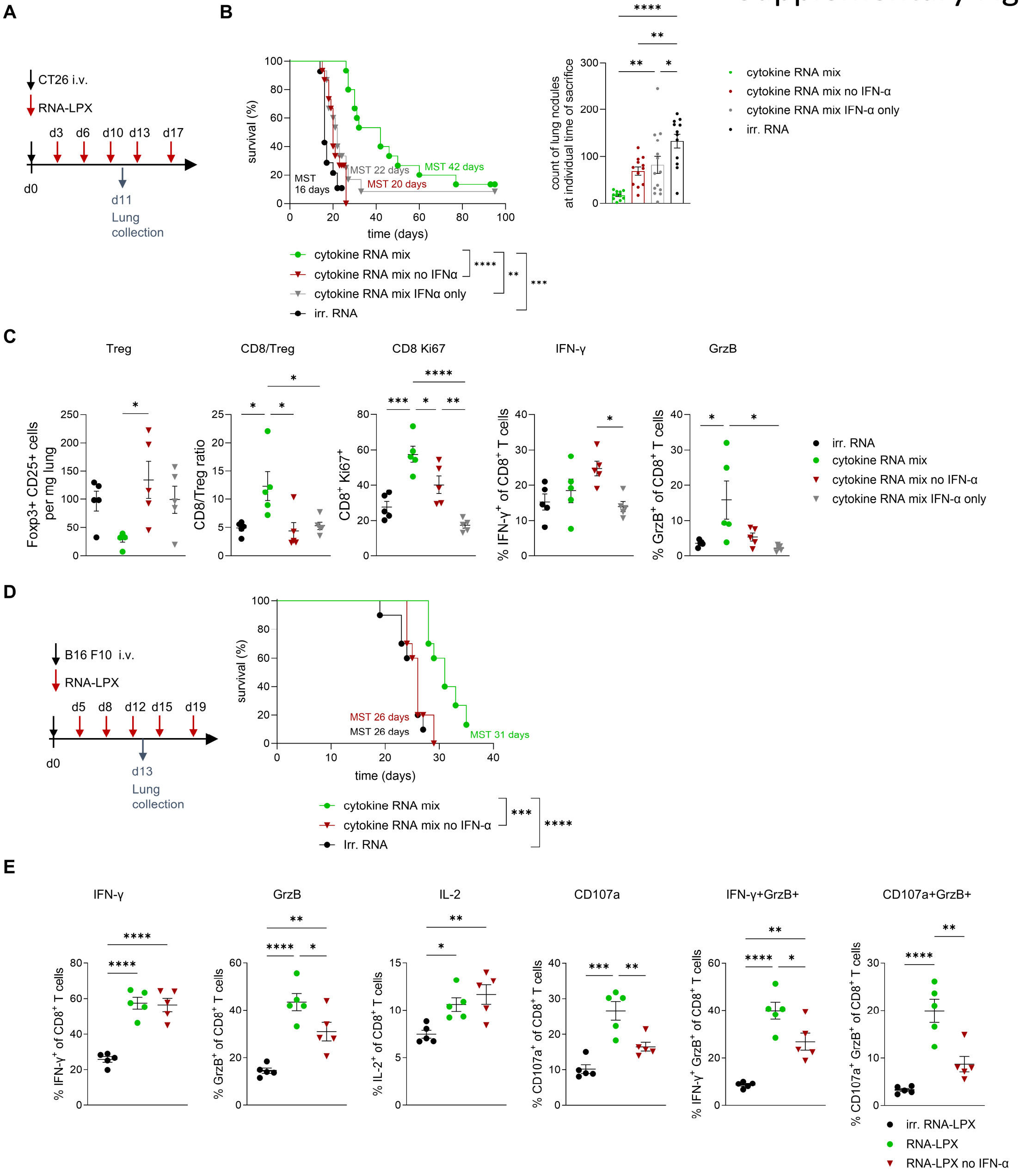


**Supplementary Figure S9. Efficacy of the cytokine RNA mix is depending on synergism of IFN-α with Alb-IL-2mut and IL-7-Alb.** (**A**) Experimental layout. Briefly: BALB/c mice (n = 15 per group for survival n = 5 for FACS analysis) were injected with CT26 tumor cells i.v., cytokine RNA mix encoding the full mix of Alb-IL-2mut (13.33 µg RNA), IFN-α (3.33 µg RNA) and IL-7-Alb (13.33 µg RNA, full mix total of 30µg RNA) or Alb-IL-2mut and IL-7-Alb (RNA-LPX no IFN-α, 13.33 µg each, total of 26.66 µg RNA), or IFN-α (RNA IFN-α, 3.33 µg) or irrelevant RNA (total 30µg RNA) as control were administered twice per week for a total of 5 injections as indicated. (**B**) Survival according to termination criteria (Mantel-Cox logrank test). Lung tumor nodule counts at the individual time point of sacrifice, significance was determined using an ordinary one-way ANOVA with post-hoc Tukey for multiple comparisons. (**C**) n=5 mice were sacrificed for lung isolation at d11 for flow cytometric analysis of tumor-bearing lungs and an ex vivo stimulation assay. Significance was determined using an ordinary one-way ANOVA with post-hoc Tukey for multiple comparisons. (**D**) Experimental layout: C57BL/6 mice (n = 10 for survival and n = 5 for FACS analysis) were injected with B16F10 tumor cells i.v. and treated with cytokine RNA mix encoding the full mix of Alb-IL-2mut, IFN-α and IL-7-Alb or Alb-IL-2mut and IL-7-Alb (RNA-LPX no IFN-α). Shown is survival according to termination criteria (Mantel-Cox logrank test). (**E**) Flow cytometric analysis of tumor-bearing lungs as indicated in (**D**), data was collected from the same mice as Fig. 2B/D and n = 5 additional mice treated with Alb-IL-2mut and IL-7-Alb (cytokine mix no IFN-a) were included for an ex vivo stimulation assay. Significance was determined using an ordinary one-way ANOVA with post-hoc Tukey for multiple comparisons.

### Supplementary Figure 10

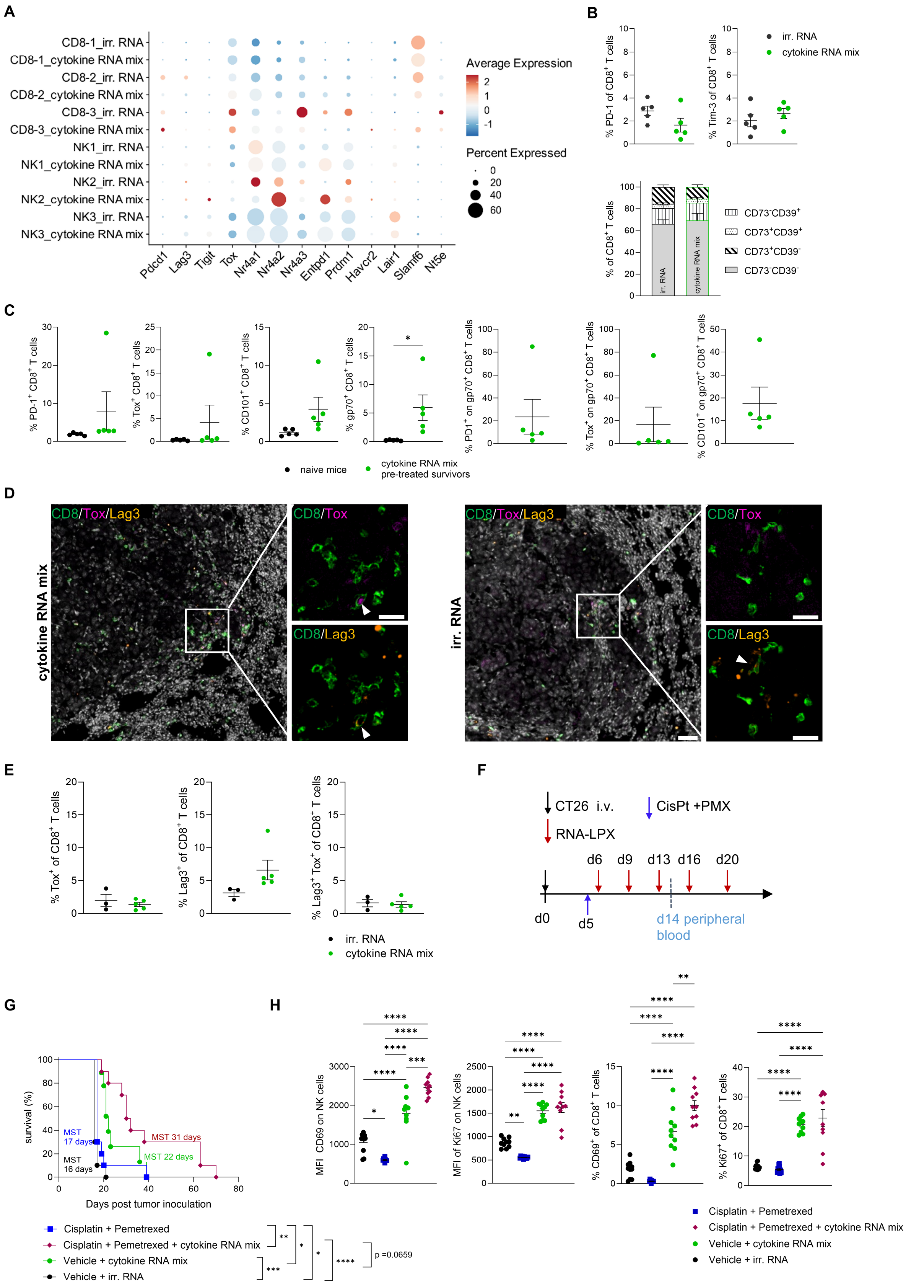

**Supplementary Figure S10. Cytokine RNA mix treatment does not majorly increase exhaustion and expression of co-inhibitory molecules and synergizes with chemotherapy.** (**A**) BALB/c mice (n=5 per group) were i.v. injected with CT26 tumor cells, cytokine RNA mix or irrelevant RNA was i.v. administered as LPX at d3, d6 and d10 p.t.i., mice were sacrificed at d11, lungs were collected and pooled at same ratios before sorting of $CD45^+$ cells. $CD45^+$ cells were subjected to scRNAseq (related to Fig. 4). Exhaustion and co-inhibitory molecule expression based on scRNAseq analyses shown as bubble plot for cytokine RNA mix-treated vs. irrelevant RNA- treated samples. (**B**) BALB/c mice (n=5 per group) were i.v. injected with CT26 tumor cells, RNA-LPX were i.v. administered at d3, d6 and d10 p.t.i., mice were sacrificed at d11, lungs were collected and analyzed via flow cytometry. Expression of PD-1 and Tim-3 as well as CD73/CD39 on $CD8^+$ T cells was analyzed. Significance for PD-1 and Tim-3 was determined by unpaired two-tailed t-test, for comparison of multiple groups (CD73/CD39) two-way ANOVA with posthoc Sidak's multiple comparisons test was performed, no significant regulation of PD-1 nor Tim-3, nor among the displayed CD73/CD39 populations can be detected. (**C**) Experimental layout as shown and described in Fig. 3A (CT26 tumor model), survivor mice of Fig. 3C (n=5) were sacrificed on d41 after re-challenge (d139 after initial tumor inoculation) as well as n=5 naive mice and lungs were analyzed by flow cytometry. $CD8^+$ T cells and gp70-antigen-specific $CD8^+$ T cells are shown for expression of exhaustion markers PD-1, Tox, CD101. (**D, E**) Experimental layout as shown and described in Fig. 2A, briefly irrelevant RNA or cytokine RNA mix formulated as RNA-LPX was i.v. administered into B16-F10 tumor-bearing C57BL/6 mice 2 days after tumor inoculation. Cytokine RNA mix (n=5) and irrelevant RNA –treated mice (n=3) were sacrificed at the individual end point (cytokine RNA mix between d29-d32 and irrelevant RNA between d29-d30) and subjected to Phenocycler Fusion immunofluorescence staining. (**D**) Representative Phenocycler Fusion (CD8, green; Tox, pink; Lag3, orange; DAPI, grey). Tox and Lag3 expressing T cells are shown with white arrowheads. The majority of T cells do not express Lag3 and Tox in both groups. Scale bars: 50 µm and 20 µm. (**E**) Quantification of $Tox^+$ and/or $Lag3^+$ $CD8^+$ T cells from PhenoCycler Fusion immunofluorescence staining in cytokine RNA mix (n=5 mice) and irrelevant RNA (n=3) –treated mice. (B, C, E) Significance for pairwise comparisons was determined by unpaired two-tailed t-test (n.s.) (**F**) Experimental layout for (G, H), briefly: BALB/c mice were injected with CT26 tumor cells i.v., cytokine RNA mix or irrelevant RNA as LPX were administered twice per week for a total of 5 injections as indicated and monitored until termination criteria were reached (n = 10 per group except for Vehicle + cytokine RNA mix where 1 mouse is excluded from survival analysis due to suboptimal tumor injection). Pemetrexed (200 mg/kg) and Cisplatin (6 mg/kg) was injected one on d5. (**G**) Survival according to termination criteria, significance as calculated by Mantel-Cox logrank test. (**H**) Analysis of CD8+ T cells and NK cells in peripheral blood of the mice (n = 10 per group, MFI: median fluorescence intensity). Significance was determined using an ordinary one-way ANOVA with post-hoc Tukey for multiple comparisons.
